# Lack of the glycine receptor alpha 2 increases striatal activity and motivated behavior

**DOI:** 10.1101/2022.08.31.506020

**Authors:** Jens Devoght, Joris Comhair, Giovanni Morelli, Jean-Michel Rigo, Rudi D’Hooge, Chadi Touma, Rupert Palme, Ilse Dewachter, Martin vandeVen, Robert J. Harvey, Serge Schiffmann, Elisabeth Piccart, Bert Brône

## Abstract

Distinct developmental pathologies, including autism spectrum disorder and schizophrenia, exhibit impaired reward-motivated behavior. Key to proper reward-motivated behavior is dopamine-mediated modulation of striatal activity. The glycine alpha 2 receptor (GlyRα2) is the single functionally expressed glycine receptor in adult striatum, and is therefore ideally positioned to modulate striatal behavior and cellular activity. Here, we report excessive appetitive conditioning in male GlyRα2 knockout mice. We next show that depletion of GlyRα2 enhances dopamine-induced increases in the activity of putative dopamine D1-expressing striatal projection neurons, while not affecting dopamine neuron activity. Moreover, we found that excessive locomotor responses to amphetamine in GlyRα2 KO mice correlate with immediate early gene c-fos expression in the dorsal striatum. 3-D modeling revealed an increase in the number of activated cell ensembles in the striatum in response to D-amphetamine in GlyRα2 KO mice. Taken together, we show that depletion of GlyRα2 impairs reward-motivated behavior and altered striatal signal integration. This sheds important light onto the cellular mechanisms that underlie reward function, and pave the way towards novel therapeutics for the treatment of e.g. schizophrenia and addiction.

**Significance statement:** The glycine receptor alpha 2 has long been studied for its role in development, with expression assumed to decline throughout adulthood in favor of the glycine receptor alpha 1 and 3. Yet, we showed that in the dorsal striatum, the glycine alpha 2 receptor is the only functionally expressed glycine receptor at adult age (Molchanova et al., 2017). **In the present work, we show for the first time that the glycine alpha 2 receptor crucially affects striatal cell activity, which lies at the basis of reward-motivated behaviors, and which is impaired in many psychiatric pathologies**. Indeed, a link between the mutations in the glycine alpha 2 receptor and autism as well as schizophrenia has been described, but a functional role for the glycine alpha 2 receptor in adult brain structures that are involved in psychiatric pathologies, was never shown before.

## Introduction

The brain reward circuitry includes projections to the striatum that originate in the thalamus and cortex on the one hand, and the midbrain on the other hand [1, 2]. The striatum integrates these inputs, so providing the core cellular mechanism of reward-motivated behavior [3]. Indeed, impeded striatal development and/or signaling together with deficits in reward-related behavior are described in distinct pathologies, ranging from autism spectrum disorder (ASD) to addiction and psychosis [4–6].

Striatal projection neurons make up the main cell type within the striatum (95%). These cells receive dopaminergic inputs from the midbrain [7, 8], and can be divided into cells that expresses the Gs-coupled D1 receptor (D1R)- and cells that express the Gi-coupled D2 receptor (D2R), although co-expression is reported as well [9–11]. At rest, all SPNs reside in a hyperpolarized resting state (around -80mV), i.e. the downstate, largely governed by inwardly rectifying potassium currents [12–15]. In addition to dopaminergic input, these cells also receive glutamatergic inputs from the cortex. In case of sufficient, converging glutamatergic innervation, SPNs transition to a near-threshold upstate. Inwardly rectifying potassium currents cease and L-type calcium channels start to open. In response to motivationally relevant cues, such as for instance a reward-predicting cue, dopamine is released from the midbrain to the striatum. D1R activation increases excitability by negative modulation of K_v_1.2 channels, as well as small-conductance and big conductance potassium channels (SK and BK channels) [16, 17]. D2R activation in upstate SPNs, however, produces opposite effects: α-amino-3-hydroxy-5-methyl-4-isoxazolepropionic acid receptor (AMPAR) currents decrease, intracellular calcium is mobilized, which causes a negative modulation of calcium Cav1.3 channels, potassium currents increase and opening of voltage-gated sodium channels decreases [18, 19]. In hyperdopaminergic pathologies such as schizophrenia, increased dopamine inputs cause excessive responses to glutamatergic inputs [20].

Here, we investigated the potential of the glycine receptor alpha 2 (GlyRα2) to limit dopamine-induced increases in SPN excitability. Glycine receptors are ligand-gated ion channels that induce a fast increase in chloride conductance upon activation. They are either homo-pentamers, containing five alpha subunits, or a hetero-pentamer containing four alpha subunits and one beta subunit. There are four alpha subunits known: alpha 1-4. Until recently, the alpha 2 subunit was believed to be expressed throughout development, with expression declining towards adulthood in favor of alpha 1 and 3. However, Molchanova et al. have demonstrated that GlyRα2 remains expressed in the adult dorsal striatum. Perforated patch voltage clamp recordings showed that activation of GlyRα2 by 3mM glycine application induces a chloride current that drives the membrane potential towards the equilibrium potential of chloride (−54mV). When in downstate (i.e. -80mV), GlyRα2 activation thus depolarizes the cell. Indeed, application of GlyR antagonist strychnine reduce the holding current and both strychnine application and GlyRα2 KO hyperpolarize the cell membrane potential. However, when the membrane potential depolarizes to exceed the equilibrium potential of chloride, GlyRα2 activation causes an inhibitory chloride current and shunts the depolarization [21]. Shunting inhibition by GlyRα2s is thus expected to be most significant when dopamine inputs enhance glutamatergic inputs in an upstate D1-expressing SPN. Importantly, GlyRα2 is the only functionally expressed glycine receptor in the dorsal striatum, which is critical for motor behavior, habit formation and motivated behavior [22–26]. Indeed, 3mM glycine application induces pronounced chloride currents in SPNs from acute mouse brain slices from wildtype mice, but elicits no response in SPNs from their GlyRα2 knockout littermates [21]. These GlyRs are thus ideally positioned to alter reward circuitry function. We hypothesized that depletion of GlyRα2 further increases dopamine-boosted activity in upstate SPNs, and consequently, exaggerate basal-ganglia-orchestrated behavior.

## Material and Methods

### Animals

Animals were maintained under a 12h/12h light/dark cycle with access to food and water ad libitum. All experiments were performed on adult male littermate mice (> 12 weeks). Male littermates with hemizygous presence (WT) or knockout (GlyRα2 KO) of *Glra2* on a C57BL/6 background were used during all experiments [39, 52, 64]. For the optogenetic studies, GlyRα2 KO mice were crossbred with a DAT^IREScre^Ai32(RCL-ChR2(H134R)/EYFP) mouse strain, as described in [93] on a C57BL/6 background. This triple transgenic mouse line with DAT-dependent expression of channelrhodopsin-2 (ChR2) allowed specific optogenetic stimulation of dopamine neurons in WT and GlyRα2 KO animals. Male littermates heterozygous for DAT^IREScre^ and ChR2 allele, but hemizygous presence or absence of *Glra2* were used for experiments (respectively referred as WT and GlyRα2 KO mice in these experiments). The DAT^IREScre^Ai32(RCL-ChR2(H134R)/EYFP) mouse line was bred and provided by Prof. M. Beckstead (JAX stock numbers #006660 and #012569). Animal experiments were approved by the local ethical committee at Hasselt University and in conformity with the EU directive 2010/63/EU on the protection of animals used for scientific purposes.

### Acute brain slice preparation

Acute brain slices were prepared for electrophysiological recordings and imaging experiments. Animals were cervically dislocated and brains were rapidly isolated into oxygenated (95% O_2_ and 5% CO_2_ mixture) ice-cold cutting artificial cerebrospinal fluid (C-aCSF) containing (in mM): 140 choline chloride, 2.5 KCl, 1.25 NaH_2_PO_4_, 7 MgCl_2_, 26 NaHCO_3_, 0.5 CaCl_2_, 11.1 D-glucose. Sagittal slices (250 µm) were prepared on a vibrating microtome (VT1200S; Leica Microsystems IR GmbH, Wetzlar, Germany) and allowed a recovery of 1h at 36°C in oxygenated recovery solution (R-aCSF) containing (in mM): 127 NaCl, 2.5 KCl, 1.25 NaH_2_PO4, 3 MgCl_2_, 26 NaHCO_3_, 2 CaCl_2_, 11.1 D-glucose. For electrophysiological measurements in the SNc oxygenated C-aCSF and R-aCSF contained (in mM): 127 NaCl, 2.5 KCl, 1.2 NaH_2_PO_4_, 1.2 MgCl_2_, 2.4 CaCl2, 21.4 NaHCO_3_, 11.1 glucose and 1.25 kynurenic acid. Horizontal slices of 200 µm were prepared of the ventral mesencephalon containing the SNc.

### Electrophysiology

During recordings, slices were continuously perfused at a flow rate of 1.5-2 ml/min and maintained at a temperature of 36°C with oxygenated normal aCSF containing: 127 NaCl, 2.5 KCl, 1.25 NaH_2_PO_4_, 1 MgCl_2_, 26 NaHCO_3_, 2 CaCl_2_ and 11.1 D-glucose. Whole-cell recordings of dorsal SPNs were performed using borosilicate-glass (Hilgenberg GmbH, Malsfeld, Germany) pipettes with a resistance between 5-7 MΩ. All recordings were acquired with an EPC-9 amplifier (HEKA Elektronik GmbH, Lambrecht, Germany) and the PatchMaster interface (HEKA Elektronik). Recordings were acquired at a 20 kHz sampling interval and online filtered using a Bessel 2.9 kHz filter. Series resistances were checked before onset of whole-cell experiments and followed up regularly. If a change in series resistance exceeded 30%, the recording was discarded.

Visual and electrophysiological methods were used to identify SPNs (26, 39, 49-51) in the striatum. Effects of DA modulation on intrinsic SPN excitability were recorded in whole-cell configuration using current-clamp mode with an intracellular solution (290 mOsm) containing (in mM): 125 KOH, 3 KCl, 0.022 CaCl_2_, 10 HEPES, 0.1 EGTA, 4 MgATP, 0.5 Na_2_GTP, 5 Na_2_phosphocreatine. Methanesulfonic acid was used to adjust pH to 7.2. Measurements were performed in the presence of 10 µM GABA_A_ receptor antagonist Gabazine, 100 nm GABA_B_ receptor antagonist CGP54626, 10 µM AMPA/kainate receptor antagonist CNQX, 5 μM NMDA receptor antagonist L-689,560, 0.1 μM neuronal nicotinic acetylcholine receptor antagonist DHβE. A holding current was applied to keep SPNs at - 80 mV. The rheobase, i.e. the minimal current injection to reach action potential (AP) threshold, was determined by a current injection protocol using 10 pA current steps. AP frequency was measured at rheobase + 40 pA for 3 seconds and used as a reference baseline (BL) activity. After 20 seconds, a second rheobase + 40 pA current step was applied together with optogenetically-induced DA release, using 470 nm light pulses (0.701 mW/mm^2^) of 4 ms at 20 Hz during 1 s. Here, current injections were performed to mimic SPN upstate needed for positive activity modulation by D1Rs, while D2R-mediated SPN activity modulation is not affected by up- or downstates (24, 52). After 5 minutes, the DA-modulated intrinsic SPN activity, measured as AP frequency, was measured at a rheobase + 40 pA injection combined with optogenetic stimulated DA release. Optogenetic pulses were generated by a CoolLED pE-2 (CoolLED, Andover, United Kingdom) illumination system triggered by an EPC-9 amplifier (HEKA Elektronik) and PatchMaster interface (HEKA Elektronik).

Dopaminergic neurons were visually identified as large neurons close to the medial terminal nucleus of the accessory optic tract (53). Electrophysiological properties of DA cells, i.e. spontaneous pacemaker firing (1–5 Hz) with wide extracellular waveforms (> 1.1 ms), were used to verify cell identification (54-56). Loose cell-attached recordings were performed to measure DA cell firing frequency. Recording pipettes (3-6 MΩ) contained a sodium-HEPES-based buffer (plus 20 mM NaCl; 290 mOsm/L; pH 7.35–7.40) (57). Measurements were performed in the presence of 10 µM Gabazine, 100 nm CPG54626, 10 µM CNQX, 0.1 μM DHβE. Baseline pacemaking activity was recorded for 2 minutes. Sarcosine (500 µM) and glycine at low (30 µM) and high (1 mM) concentrations were bath perfused for 6 minutes to determine the effects of GlyR activation on pacemaking activity. Firing frequency analysis was performed on the last 2 minutes to allow glycine diffusion within the brain slice. Burst firing of DA cells was induced by NMDA iontophoresis (50 mM; pH 8.2; 500 ms pulse) using an ION-100 Iontophoresis Generator (Dagan Corporation, Minneapolis, United States) triggered by an EPC-9 amplifier (HEKA Elektronik) and PatchMaster interface (HEKA Elektronik). Iontophoretic pipettes with a resistance of 100–150 MΩ were used. D-serine (25 µM) was bath applied as a NMDA receptor co-agonist. To determine the effects of GlyRα2s on burst activity, low glycine concentrations (30 µM) and 500 µM sarcosine were bath-applied 3 minutes prior and during the experiment. Glycine-induced chloride currents were recorded in whole-cell configuration voltage using voltage-clamp mode. Recording pipettes (2.5–4 MΩ) contained (in mM): 57.5 K-gluconate, 20 NaCl, 57.5 KCl, 1.5 MgCl2, 10 HEPES, 0.025 EGTA, 2 ATP, GTP. Glycine was applied for 10 s via a SF-77B Perfusion Fast-Step System (Warner Instruments, Hamden, CT, United States). Induction of phasic activity by optogenetic stimulation was verified in loose-cell attached configuration using the previously described protocol of striatal measurements.

Firing frequency recordings were analyzed using pClamp (Molecular Devices, LLC., San Jose, USA). Series resistance protocols and glycine-elicited currents were fitted and analyzed using Igor Pro 7.0.8.1 (WaveMetrics, Inc., Portland, United States).

### Imaging

Expression of c-fos, an immediate-early gene, in the striatum was measured 2 hours after D-amphetamine (5 mg/kg; i.p.) stimulation via immunofluorescence. Acute sagittal brain slices containing the striatum of WT and GlyRα2 KO mice were made as described above. Slices were fixed overnight in 4% paraformaldehyde (PFA) at 4°C and washed with Tris-buffered saline (TBS; 0.05 M Tris and 0.9% NaCl, pH 8.4) containing 0.05% Tween. Antigen retrieval was performed by incubating slices for 15 min in a citrate buffer (10mM tri-sodium citrate, 0.05% Tween 20, pH 6.0) at 95°C. Slices were cooled down on ice for 5 minutes followed by a blocking step (10% bovine serum albumin and 0.3% Triton X-100 in TBS) for 1 hour. Slices were then incubated with primary antibody rabbit anti-c-fos (1:1000, Merck Millipore, Burlington, United States) in TBS (5% BSA, 0.3% Triton X-100) for 48h at 4°C. A secondary antibody labeled with Alexa Fluor 488 (goat, 1:500, Thermo Fisher Scientific, Waltham, United States) was applied for 90 min at room temperature. A nuclear counterstaining was performed using 4ʹ,6-diamidino-2-phenylindole (DAPI) (Thermo Fisher Scientific) in TBS for 15 min. Slices were mounted in Fluoromount-G Mounting Medium (Thermo Fisher Scientific). Control immunostainings without primary antibodies were performed and showed no non-specific signals.

Microscopy images (Z-stacks) were acquired with a Zeiss LSM510 META (Carl Zeiss Microscopy GmbH, Jena, Germany) confocal microscope system mounted on an inverted Axiovert 200 M. A Plan-Apochromat 40×/0.95 Korr (Carl Zeiss Microscopy GmbH) air objective with cover slip adjustment (0.17) was used during image acquisition. A continuous wave Argon laser (488 nm; LASOS Lasertechnik GmbH, Jena, Germany) and a MaiTai DeepSee (790 nm; Spectra-Physics Inc., Santa Clara, USA) pulsed-laser were used to excite Alexa 488 and DAPI respectively. During the measurements laser power was kept low (5 mW at sample position) to limit the amount of photobleaching while collecting Z-Stacks. BP 500-550 IR and BP 390-465 IR emission filters were used. Images were acquired using the ZEN 2009 software (Carl Zeiss Microscopy GmbH). Z-stack tissue volumes contained typically 21 slices carrying dimensions of 225×225×42 µm. Pixel dwell time was 3.2 µsec. Limitations set by photobleaching required rapid collection of 2 μm step size Z-stacks over a depth of about 40 μm.

Processing and analysis of the acquired dual channel c-fos and DAPI nuclear stain confocal image Z-stacks were carried out both with the open source GNU General Public License image processing package ImageJ FIJI [94]. and commercial Matlab R2021b (MathWorks, Eindhoven, The Netherlands). LSM Toolbox [95] and Bio-Formats plugin scripts [96] were used to read the data. DAPI stained Z-Stacks showed a very dense cell content. Specimen tissues carried a large variability dependent on treatment and control towards background and noise levels also regionally within individual optical slices. As reviewed and discussed elsewhere [97–99], these observations in combination with the increased scattering and weaker signals from deeper layers in the Z-Stacks made automatic segmentation of the c-fos channel a challenge. To minimize an undercounting bias of c-fos positive cells that can amount to 50% in the deeper tissue layers of the raw Z-stacks, due to intensity decay, images were normalized considering the histograms of the individual slices in combination with a Z-stack normalization near the mean (optical slice 11) of the Z-stack as indicated by [100]. Effects of uneven illumination were reduced via a generalized neighborhood non-local means denoising plugin [101–103]. Reduction of the influence of locally varying overall background via a rolling ball approach had limited success. Subsequently for optimization and validation during interactive morphological segmentation [104] enhanced 2D Z-Projection color images - showing the outlines of brighter c-fos objects discernible above the noisy background - were simultaneously displayed side-by-side with the Z-projections of the binary corrected Z-Stacks. Operator supervised iterative adjustment of automated thresholding proved most effective and least time-consuming. Segmentation quality was validated by expert ground truth verification. Cell counting was performed manually on the 2D Z-projections. Z-stacks were visually checked throughout all optical slices.

### RT-qPCR

For the quantification of subunit expression in the midbrain, the SNc was dissected from WT and GlyRα2KO mice and homogenized. Total RNA was isolated by the guanidinium thiocyanate-phenol-chloroform procedure using QIAzol lysis reagent (Qiagen, Hilden, Germany) followed by RNeasy Kit (Qiagen) purification. RNA purity and quantity were checked by NanoDrop 1000 Spectrophotometer (Thermo Fisher Scientific). Purified RNA was converted to cDNA using the High-Capacity cDNA Reverse Transcription Kit (Applied Biosystems, Foster City, United States). RT-qPCR was performed in a Fast SYBR Green Master Mix (Applied Biosystems) containing 12.5 ng cDNA template, 3 mM forward (GLRA1: 5’-ATCACAAGAGCCCCATGCTAAA-3’, GLRA2: 5’-CACTGGCAAGTTTACCTGCAT-3’, GLRA3: 5’-GACGGAAGCTTTTGCACTGG-3’, GLRA4: 5’-CAGCATCAGATTGACCCTCA-3’, GLRB: 5’-TGAGCTGCTGAAACTTCCGT-3’) and 3 mM reverse (GLRA1: 5’-TGTTGTTGTTGTTGGCACCC-3’, GLRA2: 5’-GGAGACCCAGGACAAAATGA-3’, GLRA3: 5’-GACAGGGCCCCATTCCATAG-3’, GLRA4: 5’-GCAGGAGCATCTTCTAGCCA-3’, GLRB: 5’-CTCGGGTGACTGCTGAGATG-3’) oligonucleotide primer (Integrated DNA Technologies, Inc., Leuven, Belgium), and RNAse-free water was added to a total reaction volume of 10 μl. Expression levels of target genes were normalized against the most stable housekeeping genes, Gadph and Hprt, determined by qBasePlus software (Biogazelle, Ghent, Belgium). Comparative threshold cycle (Ct) quantitation (59) was performed using a StepOnePlus Real-Time PCR System (Applied Biosystems). Expression levels were converted to fold changes against the WT mean value. PCR product specificity was evaluated by melting curve analysis (StepOne Software V2.3; Applied Biosystems). Primer efficiencies between 95-105% were checked using standard curves of pooled samples and equation: E = 10[–1/slope].

Relative *c-fos* gene expression level, relevant to dopamine-modulated striatal activity, was measured 2 hours after D-amphetamine (5 mg/kg; i.p.) stimulated and saline control animals. The striatum was dissected from WT and GlyRα2 KO mice and homogenized. Total RNA was isolated by the guanidinium thiocyanate-phenol-chloroform procedure using QIAzol lysis reagent (Qiagen, Hilden, Germany) followed by RNeasy Kit (Qiagen) purification. RNA purity and quantity was checked by NanoDrop 1000 Spectrophotometer (Thermo Fisher Scientific). Purified RNA was converted to cDNA using the High-Capacity cDNA Reverse Transcription Kit (Applied Biosystems, Foster City, United States). RT-qPCR was performed in a Fast SYBR Green Master Mix (Applied Biosystems) containing 12.5 ng cDNA template, 3 mM forward (5’-ACGGAGAATCCGAAGGGAAAGGAA-3’) and 3 mM reverse (5’-TCTGCAACGCAGACTTCTCGTCTT-3’) oligonucleotide primer (Integrated DNA Technologies, Inc., Leuven, Belgium), and RNAse-free water was added to a total reaction volume of 10 μl. Expression levels of target genes were normalized against the most stable housekeeping genes, *Gadph* and *Hprt*, determined by qBasePlus software (Biogazelle, Ghent, Belgium). Comparative threshold cycle (Ct) quantitation (59) was performed using a StepOnePlus Real-Time PCR System (Applied Biosystems). Expression levels were converted to fold changes against the WT mean value. PCR product specificity was evaluated by melting curve analysis (StepOne Software V2.3; Applied Biosystems). Primer efficiencies between 95-105% were checked using standard curves of pooled samples and equation: E = 10^[–1/slope]^.

### Behavior

All animals were handled daily for two weeks prior to the start of the experiments. During all tests, animals were randomized for their genotypes and/or treatment.

#### Circadian activity

Circadian locomotor activity was monitored for 24 hours in individually housed mice in standard home cages, and measured as the number of crossings of infrared beams (3/cage), which were connected to a PC-operated counter.

#### Amphetamine-stimulated activity

Locomotion activity was video recorded within an arena (49 × 49 cm area with 30 cm walls). Baseline activity was measured for 1 hour followed by a d-amphetamine injection (5 mg/kg; i.p.) to measure dopamine-stimulated locomotion for another 2 hours. Total distance travelled was processed and analyzed in 10 min bins using the Ethovision XT (Noldus Information Technology BV, Wageningen, The Netherlands) software.

#### Appetitive conditioning

Appetitive conditioning was conducted in 16 identical automated operant chambers (Coulbourn Instruments, Allentown, PA), each set in a ventilated, sound-isolated cubicle (Coulbourn Instruments, Allentown, PA). Test cages were equipped with a grid floor connected to an electric shocker, a pellet feeder, and a nose-poke operandum. Mice (*n* = 8/group, except control: *n* = 7) were placed on a food restriction schedule, and kept at 80–90% of their free-feeding weight. Mice were trained in daily trials of 30 min during which they learned to use a nose-poking device to obtain food pellets (Noyes precision pellets, Research Diets, New Brunswick, NJ). Mice received food pellets during all trials, but the reinforcement schedule was gradually increased to obtain a stable response rate. Rate of nose poking in each trial was recorded with *Graphic State* 3.0 software (Coulbourn Instruments, Allentown, PA). Training started with one continuous reinforcement with additional guaranteed pellet delivery every 120 s (CRF + 120), followed by CRF (every nose poke rewarded), fixed ratio trials (FR5, reward on every 5th nose poke; FR10, reward on every 10th nose poke), and variable ratio trials (VR10, reward on average every 10th nose poke). Once a group of animals was stably poking (defined as no effect of time using a one-way ANOVA) for three consecutive trials, the group was moved to the next reward schedule. Transition between reward schedules happened after three days of stable poking. During six extinction trials, animals no longer received pellet rewards. On the reinstatement trial, animals were placed back on a VR10 reward schedule.

#### T-maze

The T-maze was 70 cm long, and 50 cm wide. Before training mice were allowed to explore the T-maze for ten minutes on three consecutive days. Animals were placed on a food restriction schedule, and kept at 80-90% of their free-feeding weight. Animals were then tested on six consecutive days, eight trials per day. A cup with ten food pellets was placed in the rewarded arm. In the unrewarded arm, ten pellets were placed as well to avoid odor confounds, but these were not accessible to the animal. Visual cues were placed on the end of the arms. The cue that was associated with the reward was randomized over the animals, and the location of the reward (left arm or right arm), was randomized over trials within animals. At the start of the trials, animals were placed in a start box. After 5 seconds, the start box was opened by manually lifting a divider. The latency to reach the intersection (entry with center of body) was measured and used as a measure for motor activity. Once an animal left the decision box (dark grey in Figure 1E), a divider was inserted to prevent the animals from leaving the arm of choice. Time to consume the reward was scored manually, and latency to start consuming the reward once a correct choice had been made (i.e. from the time the mouse left the decision box) was used as a measure of motivation. The number of pellets consumed was recorded for each trial as well and used as a measure for the hedonic response. xtjef

**Figure 1.**
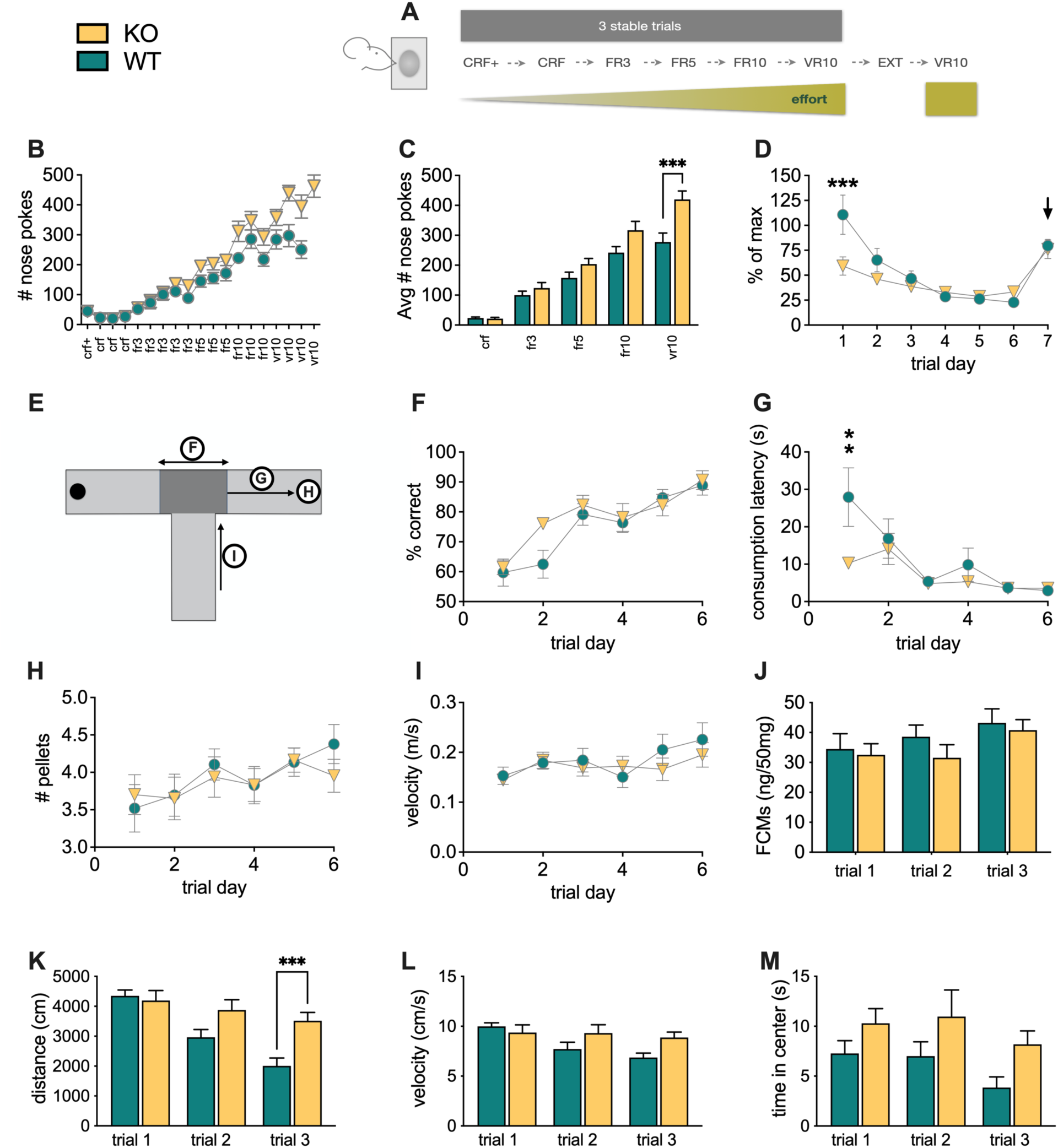
Depletion of GlyRα2 causes excessive reward-motivated behavior. (A) Graphical representation of appetitive conditioning with increasingly more demanding reward schedules. (B) GlyRα2 KO animals exhibit an increased number of nose pokes on higher demanding reward schedules, as compared to WT (crf+: continuous reinforcement + reward every 120 seconds; crf: continuous reinforcement; fr3: fixed ratio 3; fr5: fixed ratio 5; fr10: fixed ratio 10; vr10: variable ratio 10). (C) GlyRα2 KO animals exhibit enhanced responses on a vr10 reward schedule (summary bars are the average of the three days of stable nose poking within the given reward schedule). (D) GlyRα2 KO animals exhibit faster extinction rates and similar reinstatement (indicated by the arrow), compared to WT. (E) Scheme of the T-maze, indicating the data plotted in graph E-H. (F) GlyRα2 KO animals do not show enhanced associative learning compared to WT. (G) GlyRα2 KO show enhanced motivation, evidenced by the decreased time to start consuming the reward once a correct arm entry was made compared to WT. (H) GlyRα2 KO and WT littermates exhibit a similar hedonic response, evidenced by the lack of difference in the number of pellets consumed. (I) GlyRα2 KO and WT littermates approach the decision box (dark grey area in figure D) with the same velocity. Data are represented as mean ± SEM. **p<0.01, ***p<0.005. Enhanced performance in GlyRα2 knockout animals is not due to an increased stress response. (J) Levels of fecal corticosterone metabolites (FCMs) remain unaltered in GlyRα2 KO animals over consecutive trials compared to WT littermates. (K) GlyRα2 KO animals show a reduced habituation in locomotor behavior over consecutive trials compared to WT littermates. (L) GlyRα2 KO and WT animals show a similar reduction in velocity over trials. (M) GlyRα2 KO animals spend more time overall in the center of the arena, compared to WT animals. Data are represented as mean ± SEM. *p<0.05; **p<0.01; ***p<0.005

#### Anxiety and stress response

Anxiety and stress response was measured in an open field (50×50cm). Mice were placed in the arena for 1 hour and their run tracks were recorded and analyzed using the Ethovision XT (Noldus Information Technology BV) software. Eight hours after the behavioral experiment, feces were collected as described by [38]. Briefly, mice were individually housed in cages with a grid floor that allowed for feces to drop through. Filter paper was placed on the bottom to absorb urine. Mice were habituated to the grid floors the three days before testing. Fecal corticosterone metabolites were analyzed as described by Touma et al. (2004). Briefly, each fecal sample was homogenized and an aliquot of 0.05g was shaken with 1ml of 80% methanol for 30 min on a multi vortex. After centrifugation, an aliquot of the supernatant was diluted (1:10) with assay buffer and frozen at -20°C. To determine the amount of fecal corticosterone, we used a 5α-pregnane-3h,11h,21-triol-20-one EIA, which utilizes a group-specific antibody measuring steroids with a 5α-3h,11h-diol structure. Since steroid excretion is affected by the activity of the animals, we performed this experiment in a reversed light-dark schedule (6-18 lights off). However, the arena was lit from beneath. The experiment was repeated 7 and 14 days later.

### Drugs

Gabazine, CPG54626, CNQX, DHβE, cocaine hydrochloride and D-amphetamine were purchased from Tocris Bioscience (Bristol, United Kingdom). Kynurenic acid, NMDA and sarcosine were purchased from Sigma-Aldrich (Saint Louis, United States). Glycine was purchased from VWR International, LLC (Oud-Heverlee, Belgium).

### Analysis

Statistical analysis was performed using Prism 8 (GraphPad Software, San Diego, United States). Data were checked for normality and differences in variance. An overview of the performed statistical analyses can be found in supplementary Table 1.

## Results

### Depletion of GlyRα2 causes excessive reward-motivated behavior that is not mediated by an altered stress response

Adequate signal integration of dopaminergic motivational signals with cortical sensory information is key to adequate reward-motivated behavior, and the GlyRα2 is ideally positioned to affect striatal signal integration. We hypothesized that depletion of the GlyRα2 leads to excessive reward-motivated behavior. To test this, we performed an appetitive conditioning task, in which animals were trained on increasingly demanding reward schedules (acquisition), followed by an extinction and reinstatement phase (Figure 1A) in GlyRα2 KO and WT. In the acquisition phase, stable measurements were required over three consecutive days before mice proceeded to the next schedule, as described by Piccart et al. [27] to ensure similar training levels between both groups, avoiding overtraining in one group compared to the other (Figure 1B). The mean values for each reward schedule (i.e. average of performance on three stable days) were plotted (Figure 1C). A *post hoc* test showed significantly increased performance during the most demanding task in GlyRα2 KO compared to WT. In the subsequent extinction phase, GlyRα2 depletion caused a dramatic drop in number of nose pokes relative to the last conditioning trial (i.e. % of max; Figure 1D). Although overall KO and WT performed in a similar fashion, a time and interaction effect was observed. Reinstatement occurred for both genotypes in a similar manner.

Enhanced appetitive conditioning can be the result of enhanced associative learning, hedonic response, or motivation. In order to pick apart the components that might be affected by GlyRα2 depletion we performed a T-maze, as described by [28](Figure 1E). We did not observe a main genotype effect on correct arm entries, indicating no differences in associational learning (Figure 1F). We observed an overall enhanced motivation, evident in a decreased latency to consumption once the correct path arm was chosen. However, this difference disappeared over trials as animals hit a floor effect (Figure 1G). Animals consumed more pellets over trial days, an effect that was similar in WT and GlyRα2 KO (Figure 1H). Finally, WT and KO did not differ in run velocity, which similarly increased over trials (Figure 1I).

Acute stress enhances the activity of the central amygdala [29, 30]. Enhanced activity within the central amygdala in turn dramatically increases incentive, motivated behavior [31–33]. At rest, GABA inputs to the amygdala inhibit its activity, and stress-induced hyperactivity of the amygdala always coincides with the removal of inhibition [34]. Although the inhibitory control is predominantly controlled by GABAergic input, the central amygdala also expresses GlyRα2β and GlyRα3β heteromers with a minor component of GlyRα2 and GlyRα3 homomers [35][31, 32, 36, 37]. It is thus conceivable that our results in the appetitive conditioning task are, at least partly, mediated by a disinhibited stress response in the amygdala. To test this hypothesis, we performed a repeated open field experiment and measured fecal corticosterone metabolites (FCMs) [38]. We measured similar levels of corticosterone metabolites in WT and KO (Figure 1J). GlyRα2 KO show an overall increased total distance run, distance run decreases over trials in both KO and WT, and an interaction between genotype and trial is also present (Figure 1K). We found no differences in velocity between GlyRα2 KO and their WT (Figure 1L), and observed that GlyRα2 KO spent overall more time in the center of the arena over trials (Figure 1M).

Taken together, we show impaired reward-motivated behavior in GlyRα2 KO that is due to increased associative learning and motivation, but not hedonic response or stress.

### Depletion of GlyRα2 enhances the dopamine-induced increase in activity of upstate putative D1-SPNs

Reward-motivated behavior is orchestrated by signal integration within the striatum. As the GlyRα2 is the only functionally expressed receptor within the dorsal striatum, and able to shunt excessive activity, we hypothesized that depletion of GlyRα2 causes an exaggerated increase in SPN activity in response to dopamine. In order to investigate our hypothesis, we injected current (rheobase +40pA) and measured SPN excitability (“pre-DA”). Five minutes later, we optogenetically induced dopamine release to SPNs. Another five minutes later, we optogenetically stimulated again and measured SPN excitability (“post-DA”) in both WT and GlyRα2 KO (Fig 2A, B).

**Figure 2.**
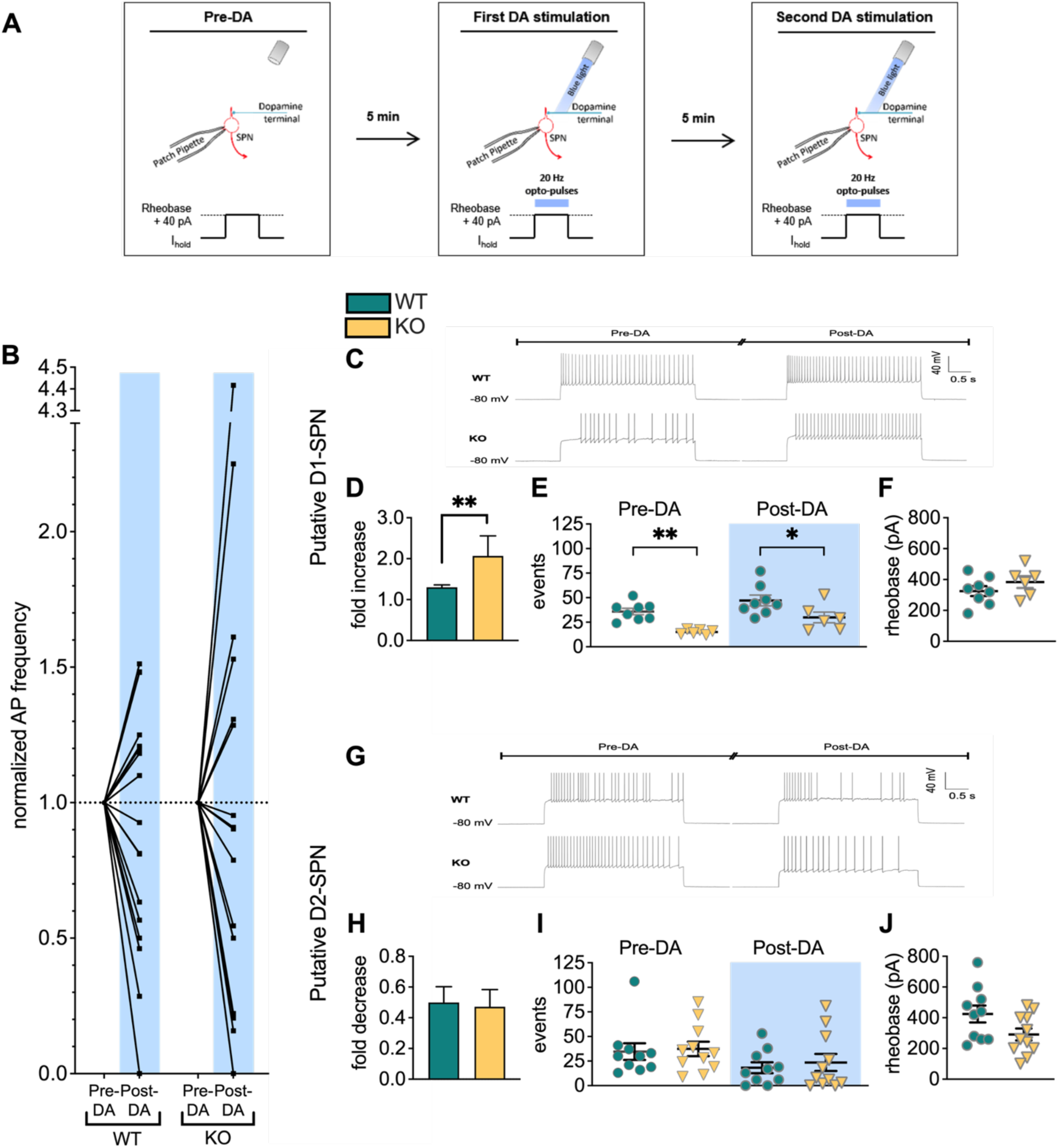
The dopamine-induced increase in activity of upstate pD1 SPNs is enhanced in GlyRα2 knockout mice. (A) Graphical protocol of dopaminergic activity modulation in SPNs. (B) Dopamine release in the striatum alters the relative intrinsic activity, measured as action potential (AP) frequency divided by baseline AP frequency, in WT (n=18) and GlyRα2 KO (n=17) mice. In both WT and GlyRα2 KO SPNs, a subpopulation of cells increased firing activity, whereas another subpopulation decreased firing activity. In agreement with distinct dopaminergic modulation of D1-versus D2-SPNs, we termed the subpopulations pD1- or D2-SPNs (pD1-SPN, pD2-SPN respectively), and analysis was conducted separately. (C) Representative traces of evoked action potentials recorded before and after dopamine modulation in pD1-SPNs of WT and GlyRα2 KO mice. (D) Dopamine-induced activity increase in pD1-SPNs is enhanced in GlyRα2 KO mice, expressed as fold change of action potential frequency after optogenetic stimulation of dopamine neurons (n_WT_ = 8; n_KO_ = 6). (E) GlyRα2 KO mice exhibit a decrease in the number of action potentials fired in pD1-SPNs, both before and after optogenetic stimulation (n_WT_ = 8; n_KO_ = 6). (F) The rheobase was not significantly different between WT and GlyRα2 KO animals in pD1-SPNs. (G) Representative traces of evoked action potentials recorded before and after dopamine modulation in pD2-SPNs of WT and GlyRα2 KO mice. (H) Dopamine-induced activity decrease in pD2-SPNs is unaltered in GlyRα2 KO mice compared to WT littermates (n_WT_ = 10; n_KO_ = 11). (I) The number of action potentials fired in pD2-SPNs is similar in WT and GlyRα2 KO mice, before and after optogenetic stimulation (n_WT_ = 10; n_KO_ = 11). See also figure S1. Data are represented as mean ± SEM. *p<0.05, **p<0.01.

We found two SPN subpopulations: one that was excited and one that was inhibited after dopamine modulation (Fig 2B). Since D1R-expressing SPNs in upstate enhance activity in response to DA, and D2R-expressing SPNs in upstate decrease activity in response to DA, (18, 19, 24, 25) we termed these cells putative D1 or D2 SPNs (pD1-SPNs and pD2-SPNs respectively) and performed further analysis in these separate populations. GlyRα2 deletion enhanced dopamine modulation of the activity of pD1-SPNs. Optogenetic stimulation increased the action potential (AP) frequency by a factor of 2.067 in GlyRα2 KO, while in WT an increase of 1.302 was observed (Fig 2C,D). However, dopamine-modulated activity of pD2-SPNs was not altered in GlyRα2 KO compared to WT (Fig 2G,H). Counterintuitively, we found that somatic current injection induced less AP firing in pD1-SPNs of GlyRα2 KO compared to WT animals. However, this difference in AP firing between genotypes became smaller after dopamine modulation, which explains the relative increase of firing pre- and post-dopamine modulation in GlyRα2 KO animals (Fig 2E,D). In pD2-SPNs, AP firing did not differ significantly between GlyRα2 KO and WT (Fig 2I). GlyRα2 KO showed no differences in rheobase (Fig 2F,J) or membrane resistance (supplementary Table 2) of pD1- or pD2-SPNs when compared to WT controls. To verify that the current injection by itself would not affect intrinsic firing, we included SPN recordings of optogenetic-inducible WT with a similar current injection protocol, but without the optogenetic stimulation. We did not observe any significant deviations in firing rates at the beginning and end of the protocol (Suppl. Fig 1B). Taken together, we show enhanced dopaminergic modulation of pD1-SPN activity in GlyRα2 KO, and reduced intrinsic excitability.

### Depletion of GlyRα2 does not affect dopaminergic neuron activity

SPN output is dependent on the glutamatergic input from the cortex, on dopaminergic input from the midbrain and on integration of both inputs in SPNs. We have previously shown reduced functional cortical input due to impaired maturation of silent synapses in GlyRα2 KO mice [39]. In the experiments described above, however, we circumvent glutamatergic input by bringing SPNs into an upstate using voltage-clamp. In the experiments described above, dopamine release was induced using optogenetic stimulation. A switch occurs from homomeric GlyRα2 in neonatal to heteromeric α1/β in adult dopamine neurons of the substantia nigra, similar to what has been described for the spinal cord and brainstem. This is supported by the relatively high occurrence of high-conductance single channel patch clamp recordings which then greatly reduce after postnatal day nineteen. Moreover, application of picrotoxin, which selectively blocks homomeric glycine receptors, strongly reduces glycine currents neonatally, but is much less efficient at doing so after postnatal day 19 [40]. It is not clear whether remaining GlyRα2 signaling at adult age alters dopaminergic activity. Therefore, we first determined the GlyR subunit expression in the substania nigra pars compacta (SNc) profile using real-time polymerase chain reaction (RT-PCR) (Figure 3A). As expected, GlyRα2 KO exhibit a complete loss of GlyR α2 subunit mRNA, with no changes in the expression of other GlyR subunit genes.

**Figure 3.**
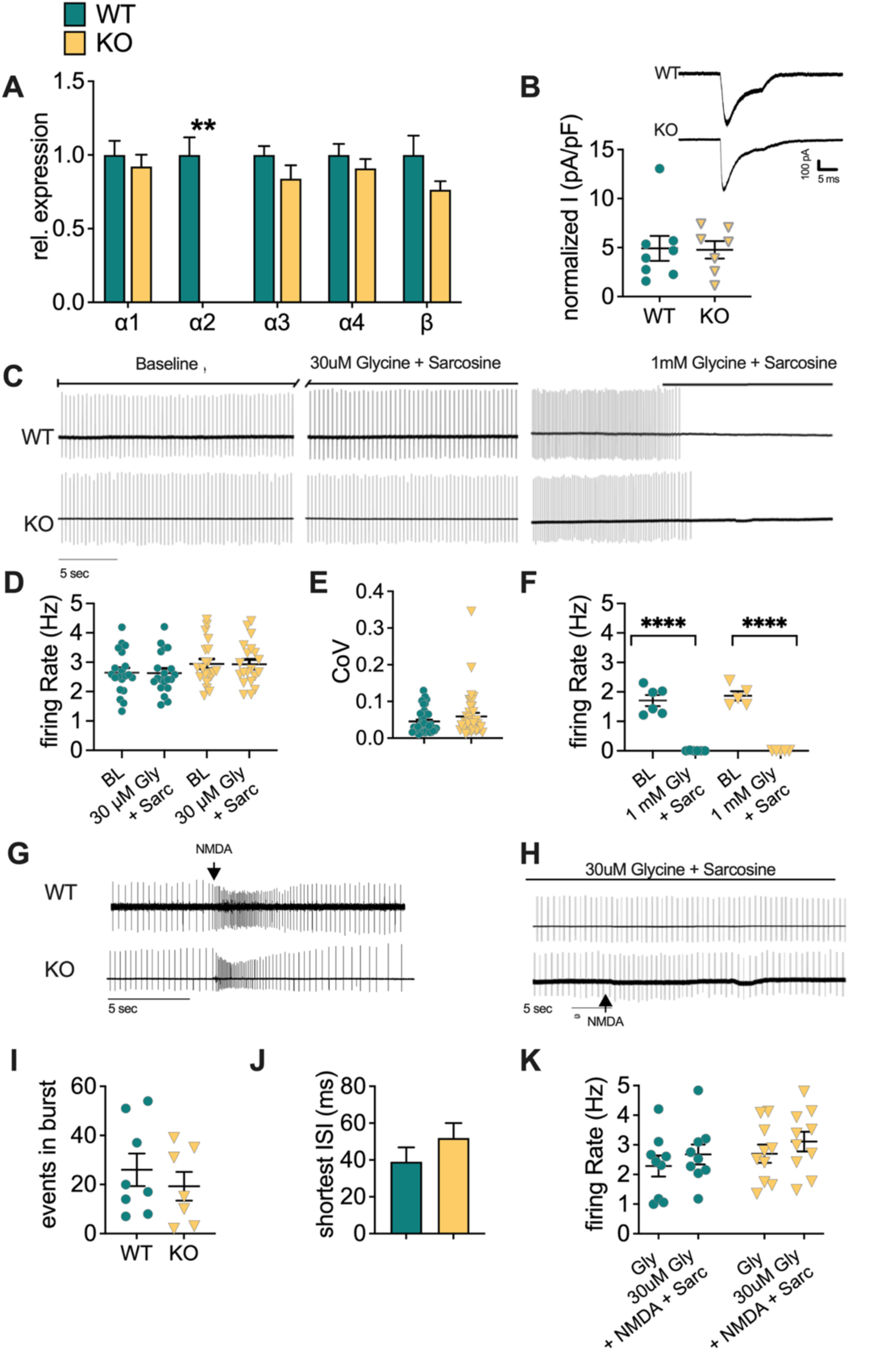
Depletion of GlyRα2 does not affect dopaminergic neuron activity. (A) Relative expression of GlyR subunits is unaltered between GlyRα2 KO and WT littermates, with the exception of α2. (B) Normalized current density is unaltered in GlyRα2 KO animals (inset: exemplar traces of glycine-induced currents in dopamine neurons from WT (top) and GlyRα2 KO (bottom) animals). (C) Exemplar traces of pacemaking firing without glycine application (left), with 30 µM glycine and sarcosine application (middle), and with 1 mM glycine and sarcosine application. (D) The baseline firing rate is unaltered in GlyRα2 KO animals, compared to WT littermates. Moreover, there is no change in baseline firing rate in response to 30 µM glycine and sarcosine in GlyRα2 KO nor WT mice. (E) Dopamine cells from GlyRα2 and WT littermates fire equally regularly. (F) Glycine (1 mM) and sarcosine completely abolishes cell firing in both GlyRα2 KO and WT littermates. (G) Exemplary traces of burst firing induced by NMDA iontophoresis in WT (top) and GlyRα2 KO animals (bottom). (H) Exemplar traces of the response to NMDA iontophoresis when 30 µM and sarcosine is co-applied in WT (top) and GlyRα2 KO mice (bottom). (I) The number of events in a burst is unaltered in GlyRα2 KO animals compared to WT littermates. (J) The shortest inter-spike interval in a burst is unaltered in GlyRα2 KO animals compared to WT littermates. (K) NMDA iontophoresis is unable to evoke burst firing in both GlyRα2 KO and WT animals in the presence of 30 µM glycine and sarcosine application. See also Figure S2. Data are represented as mean ± SEM. ****p<0.0001.

We next sought to determine the role of GlyRα2 in SNc dopamine neuron pacemaking firing. First, we measured pacemaking firing in SNc DA neurons in the presence of GABA_A_ receptor blockers, as GABA signaling can affect DA neuron firing. GlyRα2 KO showed no differences in baseline pacemaking activity compared to WT (Figure 3B, C, D), and dopamine neurons fired equally regularly (Figure 3E) with a similar inter-spike interval (Figure S2A). Indeed, a tonic current did not appear to cause differences in pacemaking firing, as we found no change in firing rate upon strychnine application (Figure S2D, E). To exclude indirect effects mediated by GABA_A_ receptors, we also measured the firing rate in the absence of GABA_A_R antagonists, but no difference was apparent in firing rate between cells from WT and GlyRα2 KO (Figure S2F, G). In order to investigate influences of GlyR activation on pacemaking activity, we bath-applied glycine both at a low concentration, which is thought to activate high affinity extra-synaptic GlyRs, and at a high concentration, which activates low-affinity synaptic receptors. Application of low levels of glycine (30 µM) did not alter firing rates (Figure 3C, D). However, high glycine concentrations (1 mM) inhibited pacemaking activity completely in both GlyRα2 KO and WT controls (Figure 3C, F). Moreover, with fast-application of glycine (1 mM) we were able to evoke currents in WT as well as GlyRα2 KO, without any difference (Figure 2B). These findings confirm the presence of functional GlyRs in DA cells of both genotypes, and suggest little to no contribution of GlyRα2 at these high glycine concentrations. Since tonic inhibitory currents mediated by GABA_A_ receptors can suppress burst firing (Lobb, Wilson, Paladini, 2010), it could be expected that GlyR-mediated tonic currents result in similar effects. We therefore measured burst activity induced by N-methyl-D-aspartate (NMDA) (50 mM) iontophoresis in GlyRα2 KO and WT (Figure 3G). Burst activity measured by loose cell patch clamp showed no differences in number of events per burst (Figure 3I), shortest inter-spike interval (Figure 3J), mean inter-spike interval (Figure S2B) or burst regularity (Figure S2C). In contrast to pacemaking activity, application of low glycine concentrations (30 µM) completely blocked burst firing in both GlyRα2 KO and WT mice (Figure 3H, K). These data confirm a role for GlyRs in activity modulation of dopamine neurons, but independent of the α2 subunit.

### *In vivo* increased dopamine neurotransmission enhances striatal activation and locomotor behavior in GlyRα2 KO mice

Behavioral output is mediated by a group of causally related co-active neurons, known as neuronal ensembles, and dopamine inputs to the striatum alter the size of the neuronal ensembles [41]. We wanted to assess whether. To establish a correlational relationship between *in vivo* excessive behavioral responses and neuronal ensemble sizes, we treated animals with D-amphetamine (5mg/kg) and recorded horizontal locomotor activity. We opted for this less behaviorally complex task compared to an appetitive conditioning task, as reward-related behavior comprises hedonic responses, learning, motor behavior and motivation, rendering interpretation more complex. The amphetamine administration exhibits a robust readout to study the function of the motor-reward pathway. We first measured circadian activity in a home cage, yet did not detect any significant differences in beam crossings between GlyRα2 KO and WT (Figure S3A). Next, we recorded locomotor activity in WT and GlyRα2 KO after saline (Figure 4A) or D-amphetamine (5mg/kg, Figure 5B) administration. GlyRα2 KO exhibited an enhanced activity when first placed in the novel environment compared to WT, but this difference vanished over time (Figure 4A). We report that treatment with D-amphetamine (5 mg/kg) caused an excessive locomotor response in GlyRα2 KO compared to WT (Figure 4B), which was absent in response to cocaine (20mg/kg) (Figure S3). To correlate the behavioral response to excessive striatal activation, we quantified the expression of immediate early gene (IEG) c-fos after acute saline or D-amphetamine treatment in GlyRα2 KO and WT (Figure 4C, D, E), often used as a tool to study neuronal ensembles that are activated by drug self-administration [42–45]. In addition, c-fos expression is particularly useful within the framework of integration of dopaminergic and glutamatergic inputs to the striatum: while glutamatergic inputs to the striatum can induce c-fos expression, dopamine and glutamate input to SPNs in upstate combined significantly enhance c-fos expression [46–48]. In agreement with an excessive response to dopaminergic input in SPNs, we detected an increase in the number of c-fos positive cells after D-amphetamine treatment two hours after injection, when c-fos protein expression typically peaks [49]. In response to saline administration, both genotypes showed a similar number of activated cells that is higher in GlyRα2 KO than WT animals (Figure 4E). We directly correlated distance travelled by a subject with the number of c-fos positive cells for that subject, and found a significant correlation, confirming the behavioral relevance of c-fos staining (Figure 4F). Dopamine release to the striatum can activate neuronal ensembles, and increased synchronous dopamine release can increase ensemble size. Small step angular variation of the images indeed revealed a regional alignment of the c-fos-positive cells in neuronal ensembles after amphetamine stimulation in the striatum, indicating a mechanism of co-activation at times of high dopamine release. Concurrently, histograms plotting the frequency that a distance between two cells was measured (Figure 4G) confirm an increase in number of c-fos positive cells in GlyRα2 KO, yet, the distribution (i.e. at which distances are the peaks located) are comparable between WT and KO, with the highest amplitudes situated around 12, 20 and 28 μm (raw data: supplementary Table 3).

**Figure 4.**
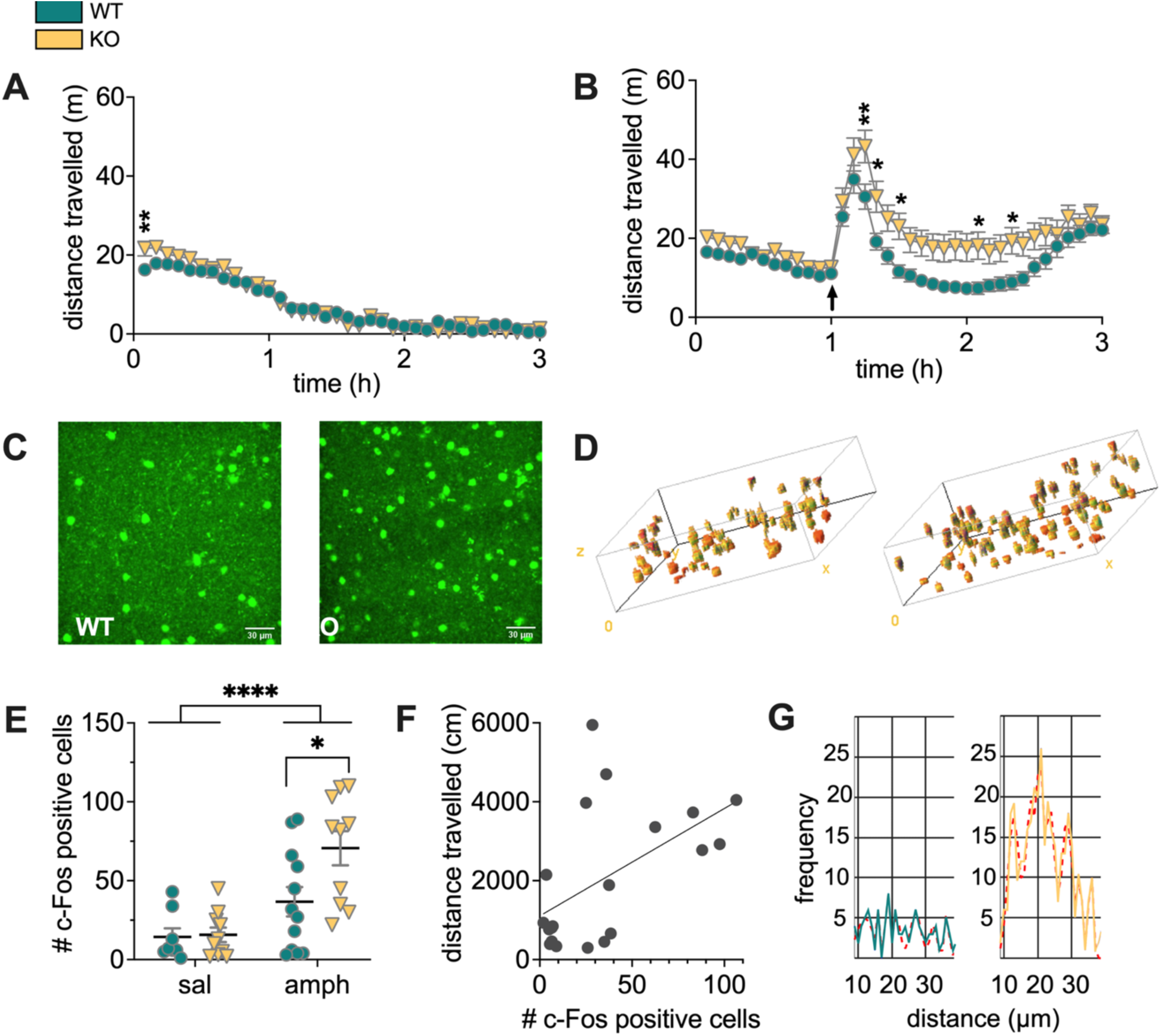
*In vivo* increased dopamine neurotransmission enhances striatal activation and locomotor behavior in GlyRα2 KO mice. (A) GlyRα2 KO animals exhibit enhanced locomotor responding in a novel environment that disappears over time. (B) GlyRα2 KO animals exhibit an increased locomotor response to 5 mg/kg D-amphetamine (i.p.). (C) Representative images of c-fos immunohistochemistry (IHC) after amphetamine (5mg/kg, i.p.) administration in WT (left) and GlyRα2 KO (right). (D) Example of 3-D processing of c-fos IHC in WT (left) and KO (right). (E) The number of c-fos positive cells increased in amphetamine-treated GlyRα2 KO animals, compared to WT littermates. (F) Distance travelled positively correlates to the number of c-fos positive cells. (G) Histograms plotting the frequency that a distance between two cells was measured indicate a similar distribution of c-fos-positive cells in WT and KO mice, suggesting similar sizes of cell ensembles (full lines: raw data, dotted red lines: Gaussian fits). Data are represented as mean ± SEM. *p<0.05; **p<0.01; ****p<0.0001.

## Discussion

The striatum plays a key role in reward-motivated behavior. Converging glutamatergic input bring SPNs to a near-threshold upstate. Concurrent dopamine release will further enhance SPN excitability. The present work investigated the potential of the GlyRα2 to affect dopaminergic modulation of SPNs in upstate, overall striatal activation, as well as striatally orchestrated behavior.

We demonstrated that amphetamine increased *in vivo* dorsal striatal SPN activity, evidenced by increased c-fos expression, that correlated to enhanced locomotor responses. Indeed, the dorsal striatum is critically involved in the motor response to psychostimulants. Dopamine efflux significantly increases in response to d-amphetamine administration [54]. Ablation of D1R-expressing SPNs in the dorso-medial striatum causes a reduction in the locomotor response to d-amphetamine [55]. Administration of cocaine causes a sharp rise in intracellular calcium levels in D1R-expressing neurons in the dorsal striatum. 3-D modeling of confocal c-fos imaging revealed an increased number of activated neuronal ensembles in GlyRα2 KO, that did not differ in ensemble size, in agreement with sparse active zone-like dopamine release sites [56]. Surprisingly, depletion of GlyRα2 enhanced the locomotor response to amphetamine, but not cocaine. This may be due to the distinct pharmacological profile of cocaine and amphetamine [57], with cocaine showing a much lower potency to induce locomotor behavior compared to amphetamine [58]. Indeed, the locomotor response to amphetamine, but not cocaine, exhibits a biphasic pattern that is typical for an enhanced response [59].

To investigate whether DA release to the dorsal SPNs increases striatal cell activity, we optogenetically stimulated DA terminals. Optogenetic stimulation appropriately mimics synchronized phasic DA release, evoking DA concentrations in the submicromolar range, similar to *in vivo* DA transients [16, 60]. Dopamine input to D1R-expressing cells in upstate enhances SPN cell excitability, whereas dopamine input to upstate D2R-expressing cells decreases cell excitability [16]. Indeed, we report that optogenetic DA release to SPNs in upstate either increases or decreases action potential frequency. We therefore termed these cells either putative D1-(pD1) or putative D2-(pD2) SPNs. It is noteworthy that Prager et al. (2020) found that DA modulates D1-SPNs in striatal striosomes and matrix in an opposite manner: DA uncaging onto striosome D1-SPNs decreased upstate duration, whereas DA uncaging onto matrix D1-SPN increased upstate duration, and this modulation depended on L-type voltage-gated calcium channels. While we cannot exclude that a fraction of the pD2-SPNs are in fact striosomal D1-SPNs, it seems unlikely given that L-VGCCs do not affect SPN excitability. In addition, the effects of dopamine on upstate duration followed a U-shape, and thus crucially depend on experimental design, and DA-induced shortening of striosomal D1-SPN upstate duration was very modest compared to the increase seen in matrix D1-SPNs [16, 61].

We next report that depletion of GlyRα2 enhanced the DA-induced increase in action potential frequency in pD1-SPNs. We note that striatal GlyRα2 are thought to be extra-synaptic receptors that produce a tonic current [21]. At low transmembrane conductances, shunting inhibition may not provide strong inhibitory effects on the cell. However, high GlyRα2 conductance can be inhibitory through shunting, thereby decreasing firing probability of the cell, similar to the inhibitory effects of high tonic GABA_A_R conductances [62]. Nonetheless, despite a larger relative increase in action potential firing frequency after optogenetic dopamine release, the absolute number of action potentials is attenuated in pD1-SPNs in GlyRα2 KO compared to pD1-SPNs from WT, both before and after dopamine release. This is in apparent contradiction with the enhanced c-fos expression. However, we must draw a distinction between activity at the population level (i.e. the neuronal ensembles as defined above) and at the level of a single cell. C-fos activity is used to study neuronal ensembles that encode associations between drug-related cues and psychostimulants (for a review, see [63]). The threshold for c-fos expression is lower than for action potential firing, and significant increases in c-fos positive cells in response to d-amphetamine in GlyRα2 KO mice is in agreement with increased striatum-orchestrated behaviors. In agreement with the present results did we report decreased firing rate in SPNs of GlyRα2 KO animals in Molchanova et al., 2017 [21]. SPNs express dopamine receptors and voltage-gated calcium (Cav1) channels in the dendrites where most cortical, glutamatergic input arrives. When in upstate, D1 receptor activation enhances the calcium currents mediated by Cav1 through a DARPP-32 signaling cascade. In GlyRα2 KO mice, the lack of glycinergic shunting inhibition, can enhance activation of voltage-gated calcium channels, and thereby consequently also enhance their inactivation, i.e. their transition into a nonconducting state. Indeed, it was earlier shown that GlyRα2 activation in the neonatal brain, where GlyRα2 activation is depolarizing, similar to an adult SPN in upstate, activates voltage-gated calcium channels and promotes calcium influx [64]. In addition, voltage-gated sodium channels might also enter a non-conductive state, further adding to the decreased excitability in GlyRα2 KO mice. The net effect is enhanced cell activation, evidenced by increased neuronal cell ensembles, enhanced sensitivity to dopaminergic modulation, but decreased single cell firing frequency. This suggests that neuronal ensemble size and the change in firing frequency, rather than the absolute frequency, dictates the behavioral response.

Dopamine neurons fired at the same rate in WT and GlyRα2 KO, and inter-spike intervals were equally regular. Low concentration glycine perfusion did not affect firing rate in either WT or GlyRα2 KO, suggesting that low affinity, extra-synaptic glycine receptors do not control basal firing. However, high concentration glycine fully inhibited pacemaking firing in both WT and GlyRα2 KO mice. It could be surmised that the expression of GlyRα1 and GlyRα3 compensate for the loss of GlyRα2, or that GlyRα2, even when present, has little effect on SNc neuronal activity. In agreement with compensation by other GlyR subtypes, we found that normalized current amplitudes in response to glycine application were similar in WT and GlyRα2 KO. We cannot exclude, however, that in spite of similar firing patterns in GlyRα2 KO and WT animals, there may be differences in dopamine release due to altered vesicle filling or release probability. Yet, based on the findings by Liu et al. (2018), increased vesicular release (probability) would likely reveal itself in a shift in inter-cell distance histograms, which was not the case. Taken together, we conclude that depletion of GlyRα2 modulates basal ganglia signaling at the level of the striatum.

The dorsal striatum has a pivotal role in motor behavior and habits. In addition, dopamine input from the substantia nigra to the dorsal striatum is also critical for motivated behavior [24, 65–68]. Viral restoration of dopamine signaling in the nigrostriatal pathway rescues operant conditioning [23], lesions to the dorsolateral striatum impair cue-motivated instrumental responding [22], motivated attraction to an incentive stimulus is strengthened upon injections with indirect dopamine agonist amphetamine into the dorsolateral striatum [25], and inhibiting neuronal activity in the dorsal striatum by microinjections of baclofen/muscimol decreases cocaine self-administration [26]. In humans, strong activation using fMRI is reported in the dorsal striatum in response to a reward-conditioned stimulus [69]. Since we revealed enhanced responses to dopaminergic input in the dorsal striatum, we hypothesized excessive reward-motivated behavior in GlyRα2 KO. We found that depletion of GlyRα2 causes excessive performance in an appetitive conditioning task. This is in agreement with the reported increase in ethanol consumption in mice lacking GlyRα2 [70] or mice that express ethanol-insensitive GlyRα2 subunits [71]. In our appetitive conditioning task, behavioral differences between GlyRα2 KO and WT controls only became apparent during highly-demanding motivational reward schedules, indicative of excessive motivated behavior. Accordingly, striatal dopamine depletion-induced impairment in appetitive conditioning only becomes apparent on highly demanding reward schedules [72], and increasing striatal activation by inhibition or depletion of phosphodiesterase 10A hinders appetitive conditioning only at highly demanding reward schedules [27, 73]. T-maze performance confirmed excessive motivated behavior. We did not detect any changes in hedonic response to reward, likely because these are mediated by hedonic hotspots in the ventral striatum where GlyRα1 and GlyRα3 are also present [75–78]. We furthermore report enhanced extinction of appetitive conditioning in GlyRα2 KO animals. This is in agreement with reports that show that increasing activity of the dorsal striatum by intra-striatal injection of a partial NMDAR agonist enhances extinction of appetitive conditioning [79]. Similarly, inactivation of the dorsolateral striatum by intra-striatal injection of sodium channel blocker bupivacaine impaired extinction [80]. Finally, it seems unlikely that our results were mediated by altered stress responses in KO, as they showed similar levels of corticosterone metabolites in a repeated open-field task and spent a larger amount of time in the center of the arena.

We thus show that GlyRα2 specifically alters reward-motivated behavior that is orchestrated by the dorsal striatum. This reward circuitry has been linked to distinct pathologies, ranging from autism spectrum disorder (ASD) to psychosis [81–85]. In agreement with a role for GlyRα2 to affect the circuitry involved in these pathologies were rare variants linked to ASD and neurodevelopmental disorders by large-scale next-generation sequencing studies [86–90]. Structure-function studies showed both loss, gain and altered function of GlyRα2 ASD mutations [89, 91]. A mutation in the GlyRα2 was furthermore linked to diagnosed schizophrenic patients in the South African Xhosa [92].

Taken together, we show that GlyRα2 can effectively inhibit dopamine-induced increases in striatal activity and reward-motivated behavior. These data provide a cellular mechanism that can contribute to the core symptoms in ASD in general.

## Declaration of interests

The authors declare no competing interests.

## Acknowledgements

This work was supported by FWO research grant 1519516N to E.P., KBF research grant to E.P, MRC Grant G0500833 to R.J.H., FWO research grant 1518419N, Saint Gillis Autism research grant to B.B., and UH research infrastructure funding I000220N. We thank Petra Bex and Rosette Beenaerts for technical assistance and Michael J. Beckstead for providing us with DAT-Cre mice.

---

**Supplemental Figure 1.**
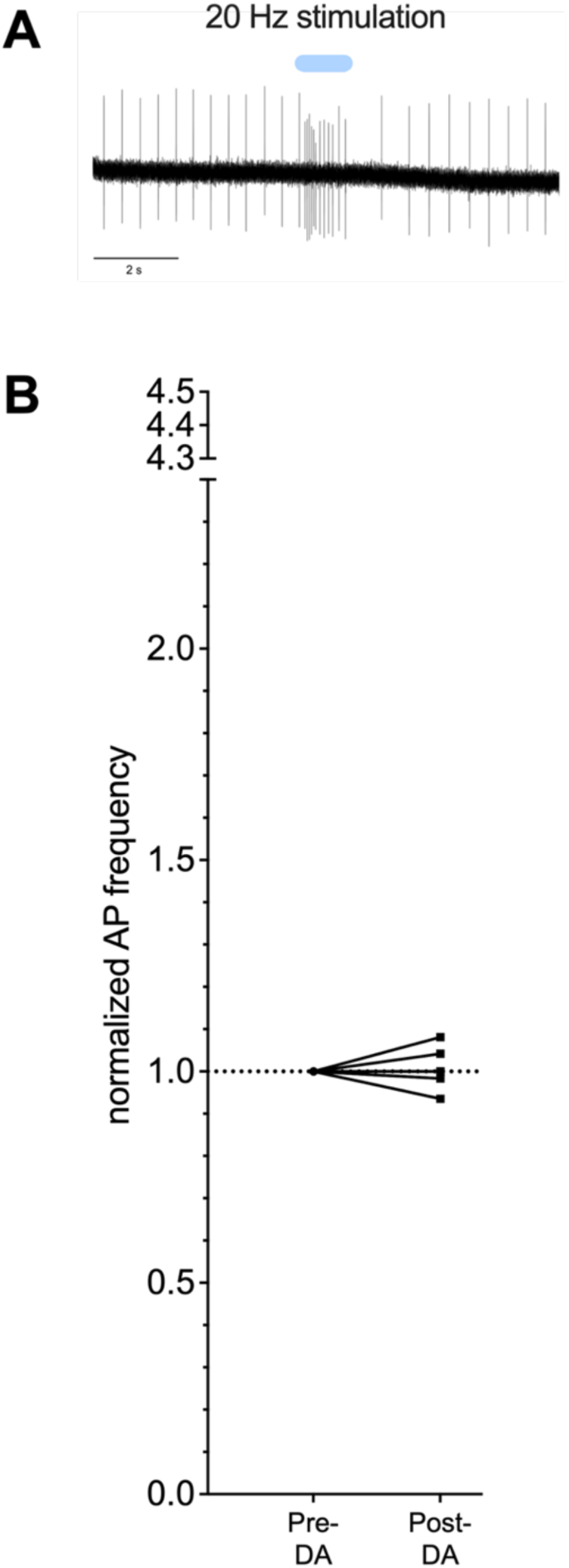
(A) Optogenetic stimulation (10 pulses, 20 Hz) of channelrhodopsin-2-expressing dopamine cells in the substantia nigra pars compacta induces burst activity. (B) Control SPN recordings in which DA terminals were not stimulated optogenetically do not show a significant change in action potential frequency over time.

**Supplemental Figure 2.**
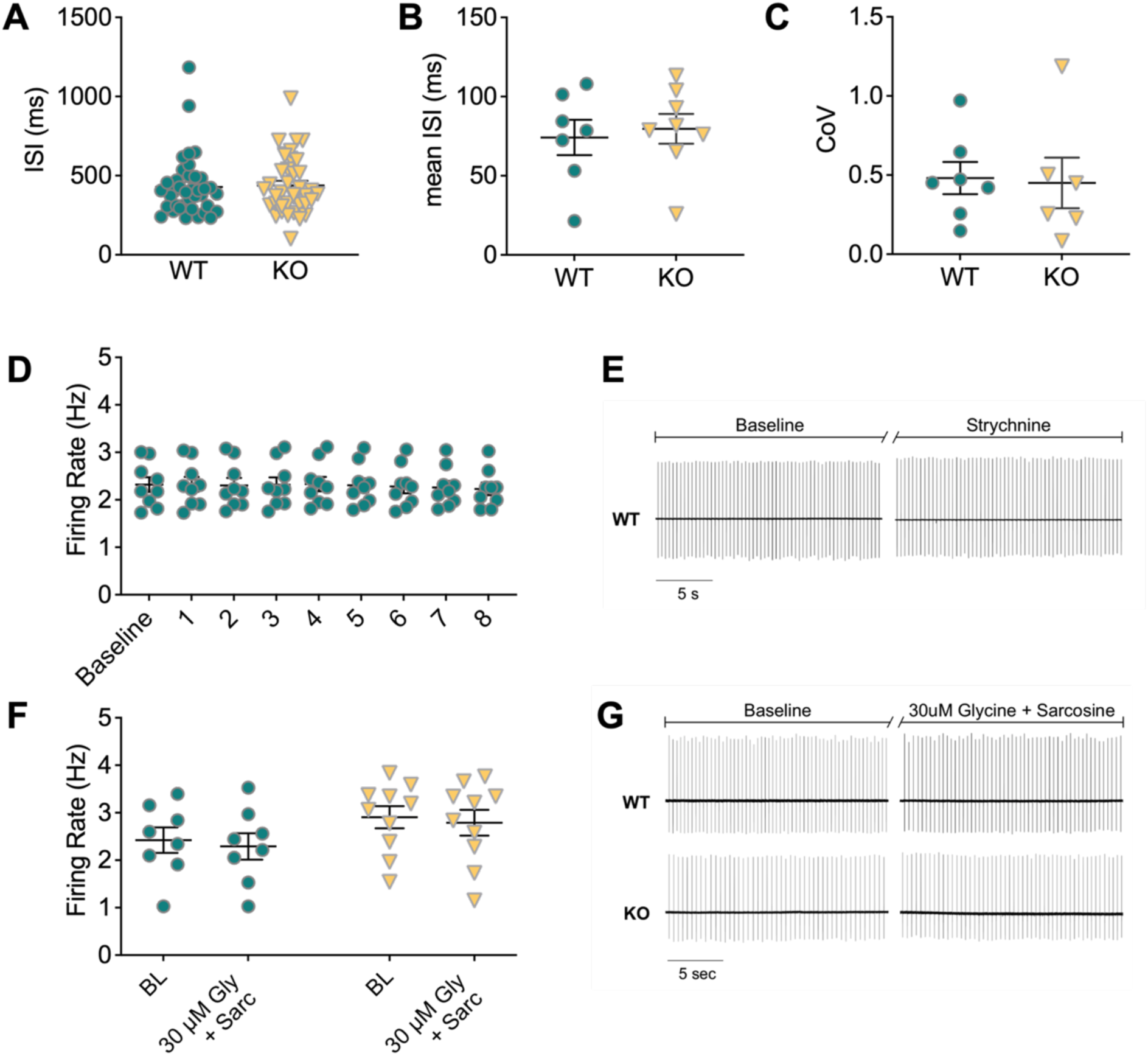
(A) Inter-spike interval during pacemaking activity of dopamine neurons is unaltered in GlyRα2 KO mice. (B) The mean inter-spike interval of a burst in dopamine cells is unaltered in GlyRα2 KOs compared to WT littermates. (C) Dopamine cells from GlyRα2 and WT littermates fire equally regularly during an NMDA-induced burst. (D) Strychnine application does not alter the pacemaking activity of dopamine cells in WT mice (represented as 1 min bins). (E) Exemplar traces of dopamine neuron pacemaking firing without (left) and with strychnine application (right) in WT mice. (F) Baseline firing rate of dopamine neurons is unaltered in GlyRα2 KO animals compared to WT littermates in the absence of GABA_A_R and GABA_B_R blockers. Moreover, there is no change in baseline firing rate in response to 30 µM glycine and sarcosine in either GlyRα2 KO or WT mice. (G) Exemplar traces of pacemaking firing in response to 30 µM glycine and sarcosine, and absence of GABA_A_R and GABA_B_R blockers in WT (top) and GlyRα2 KO mice (bottom). Data are represented as mean ± SEM.

**Supplemental Figure 3.**
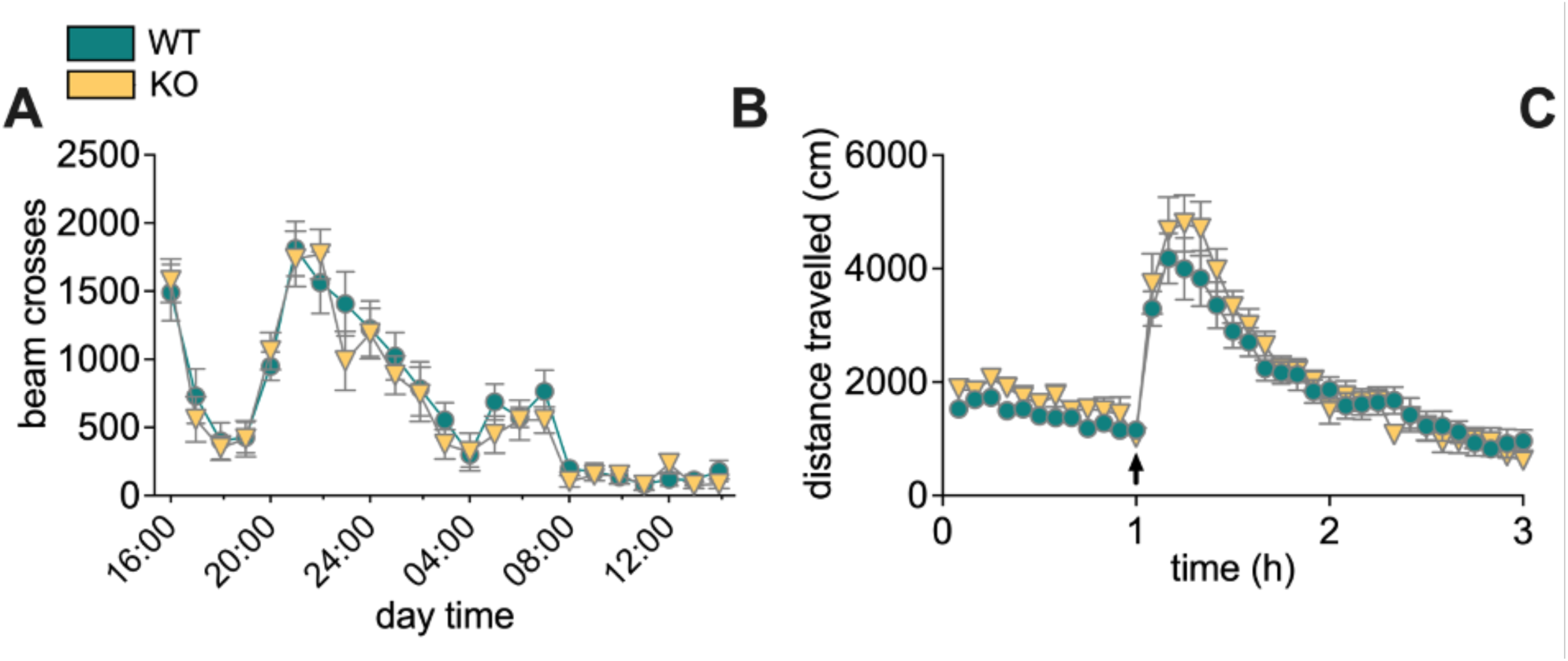
(A) GlyRα2 KO animals exhibit no changes in circadian activity compared to WT littermates. (B) The locomotor response to 20 mg/kg cocaine (i.p.) is unaltered in GlyRα2 KO animals compared to WT littermates.

**Supplemental table 1.**
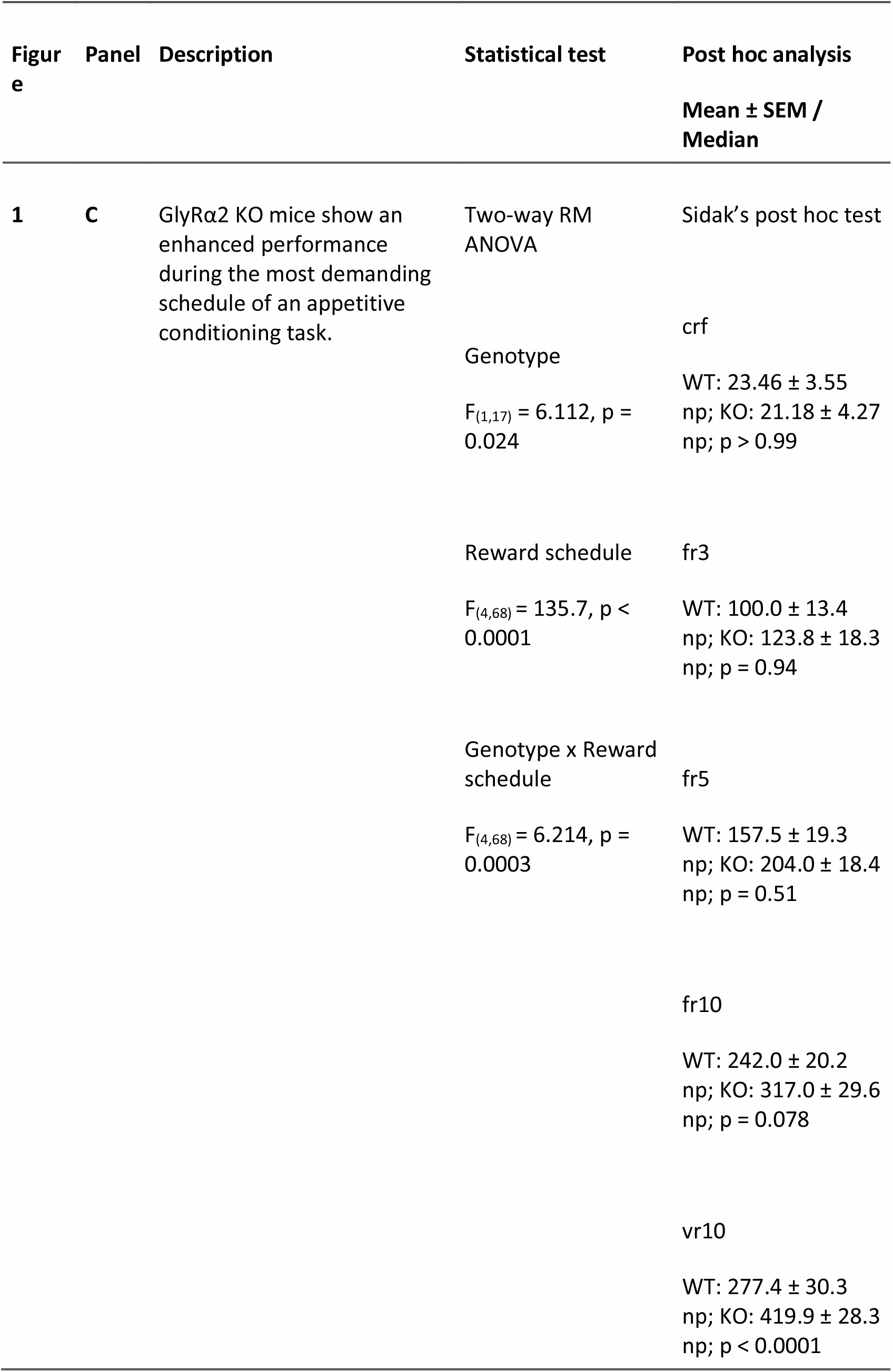
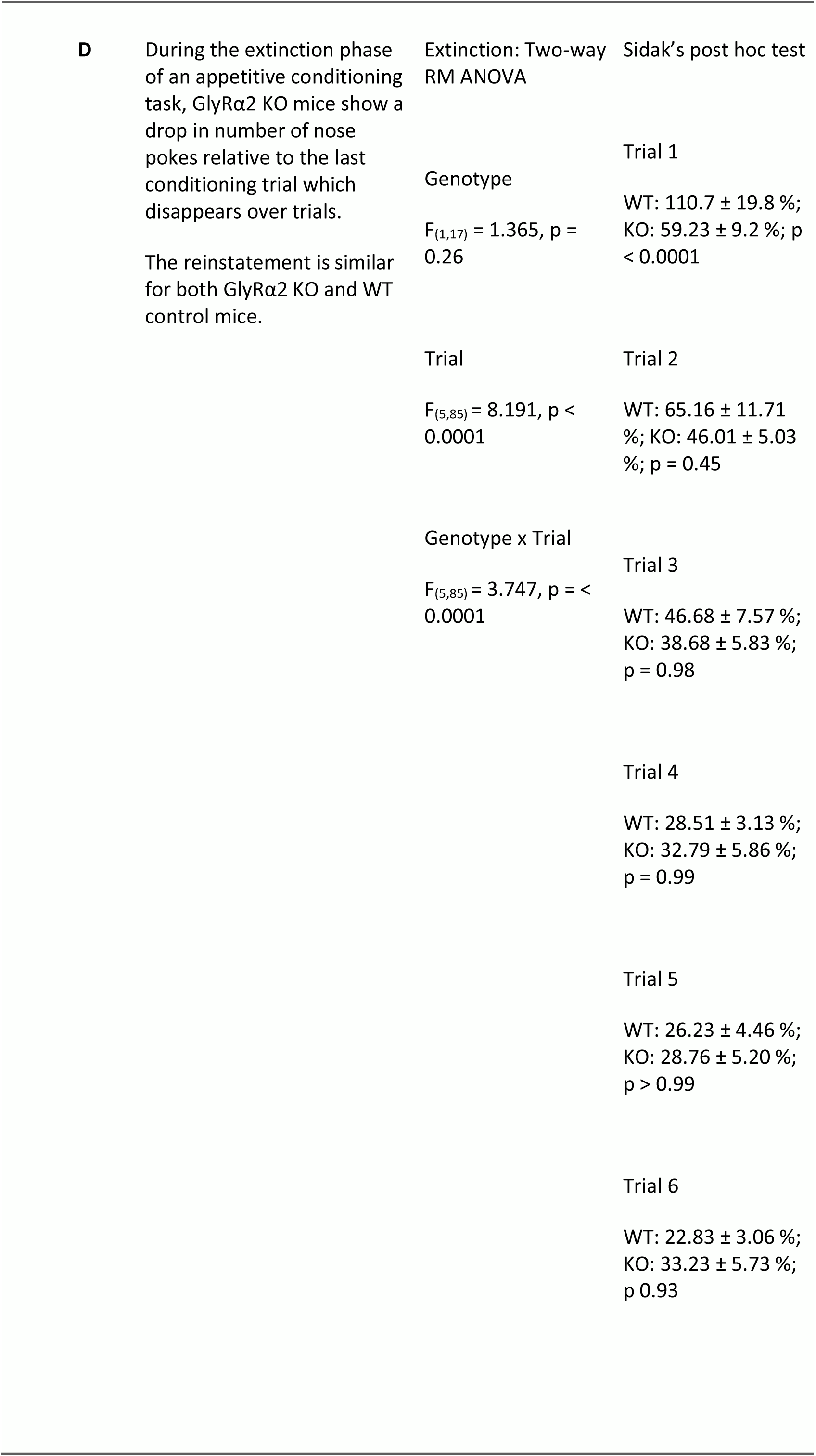
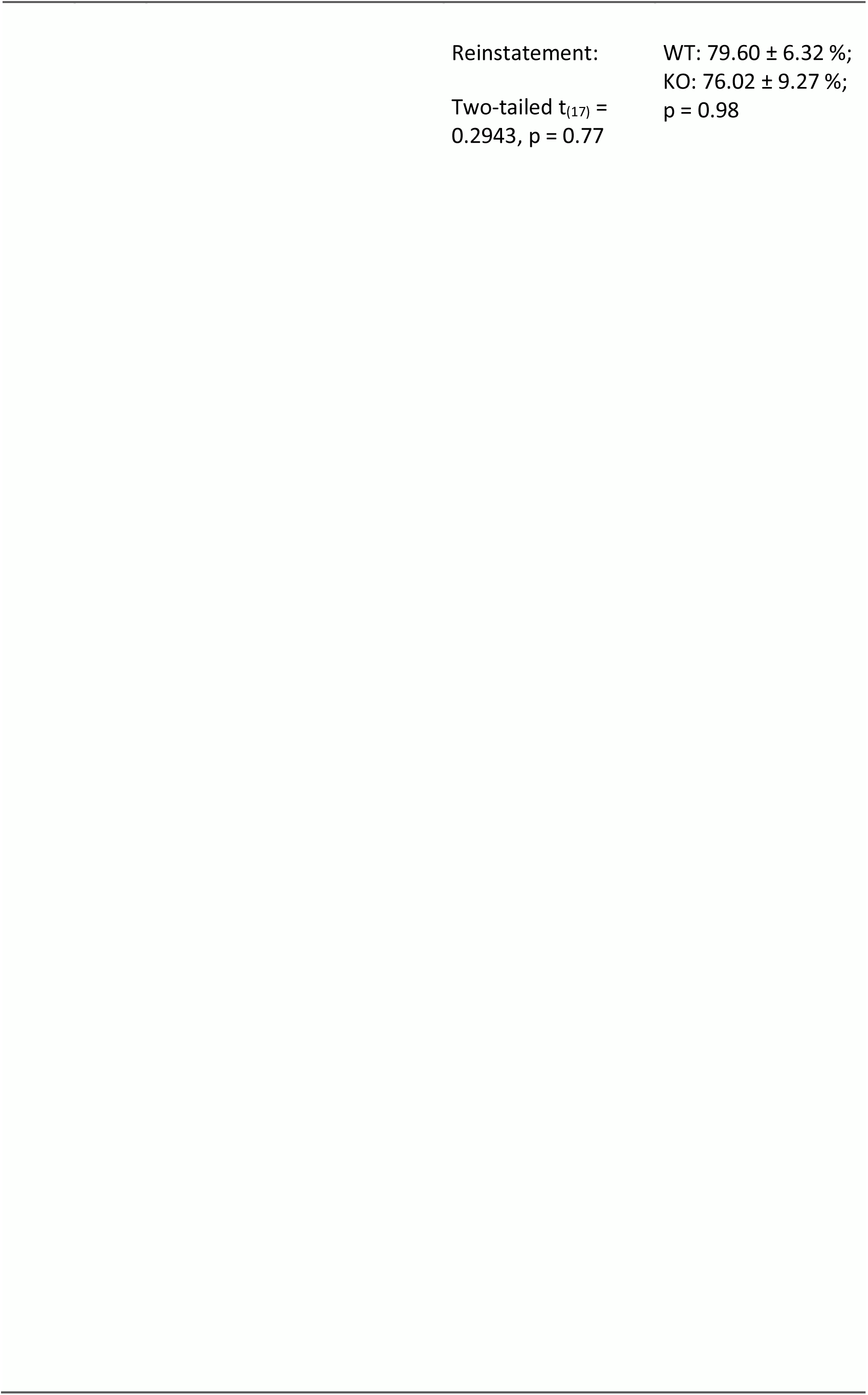
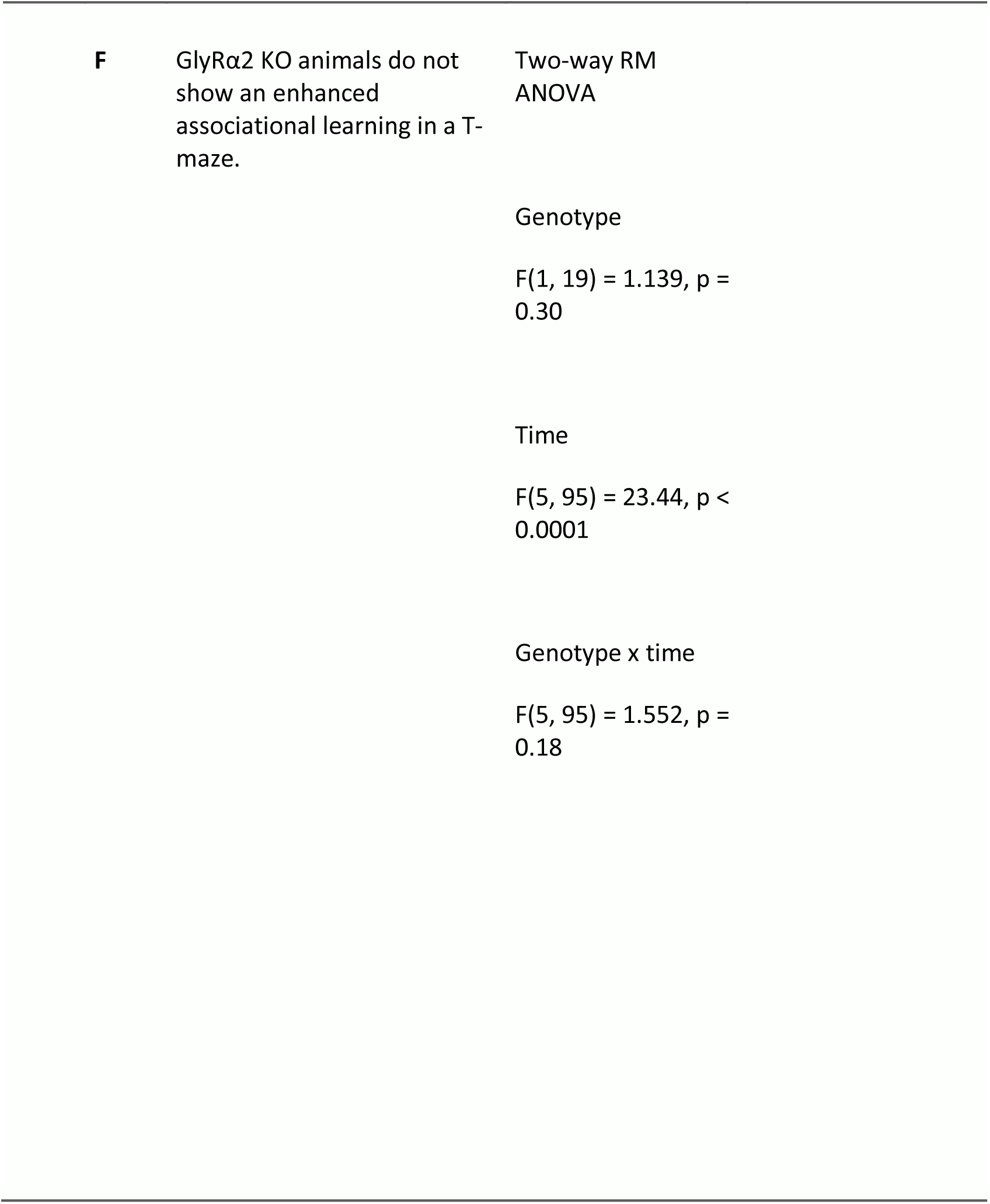
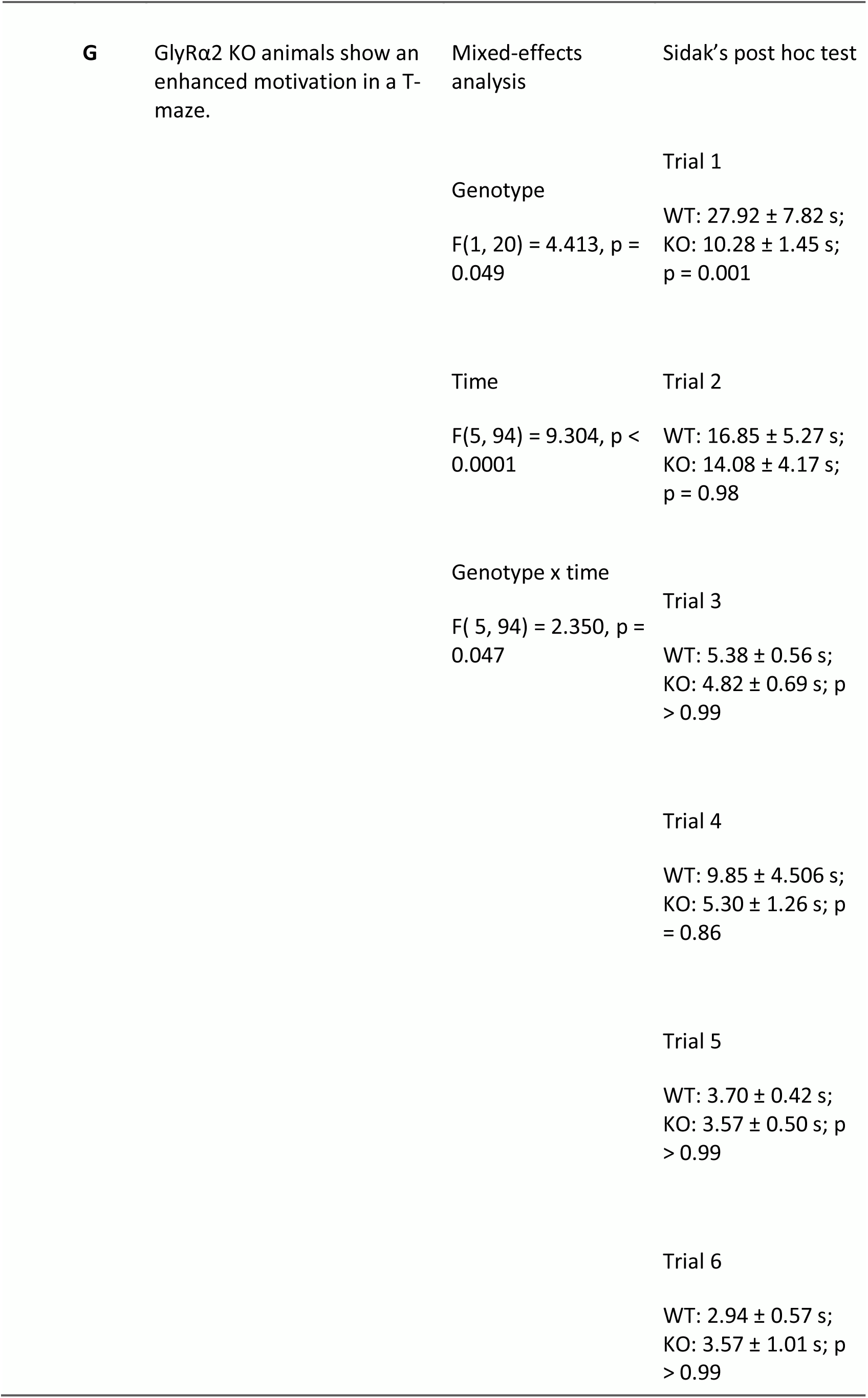
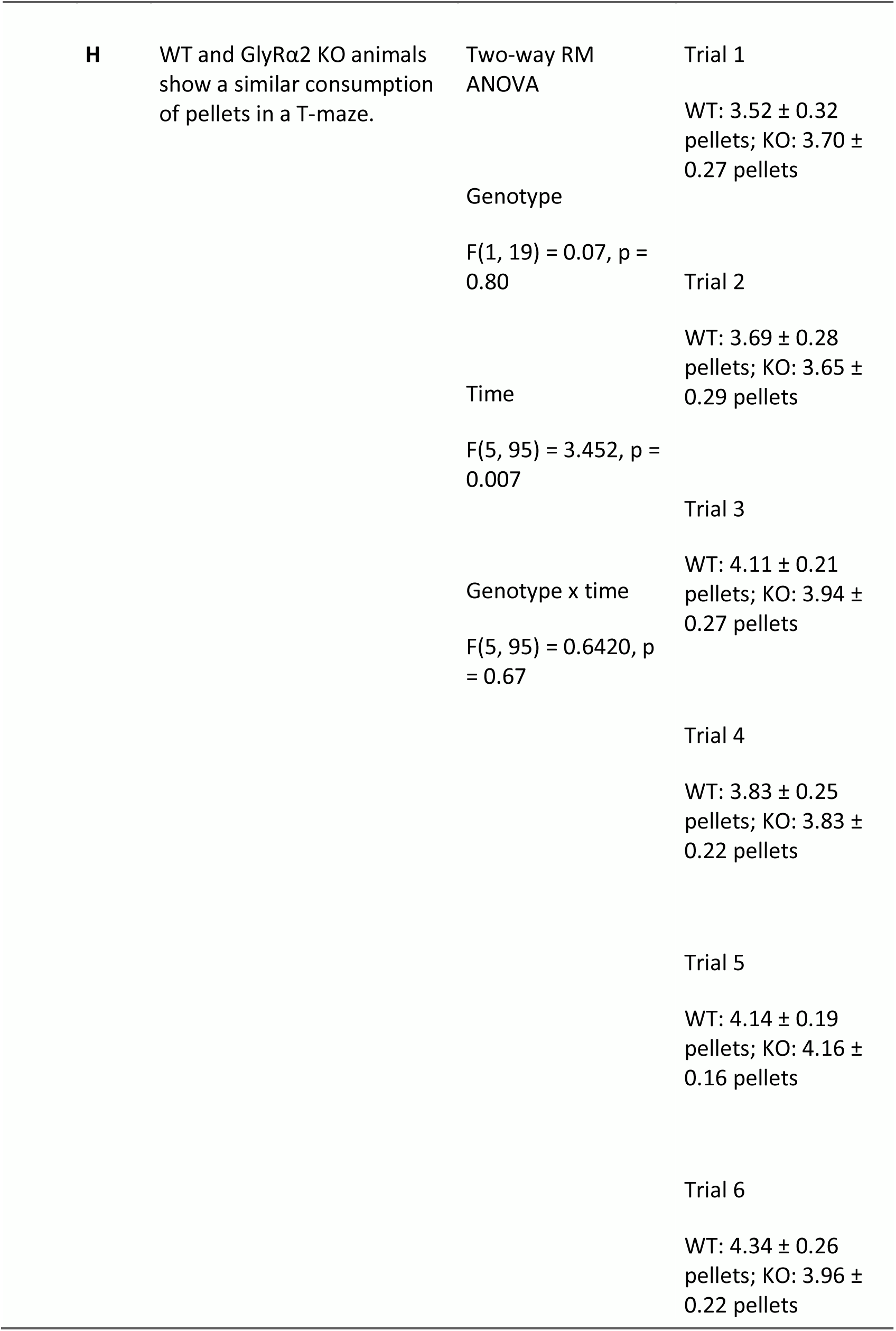
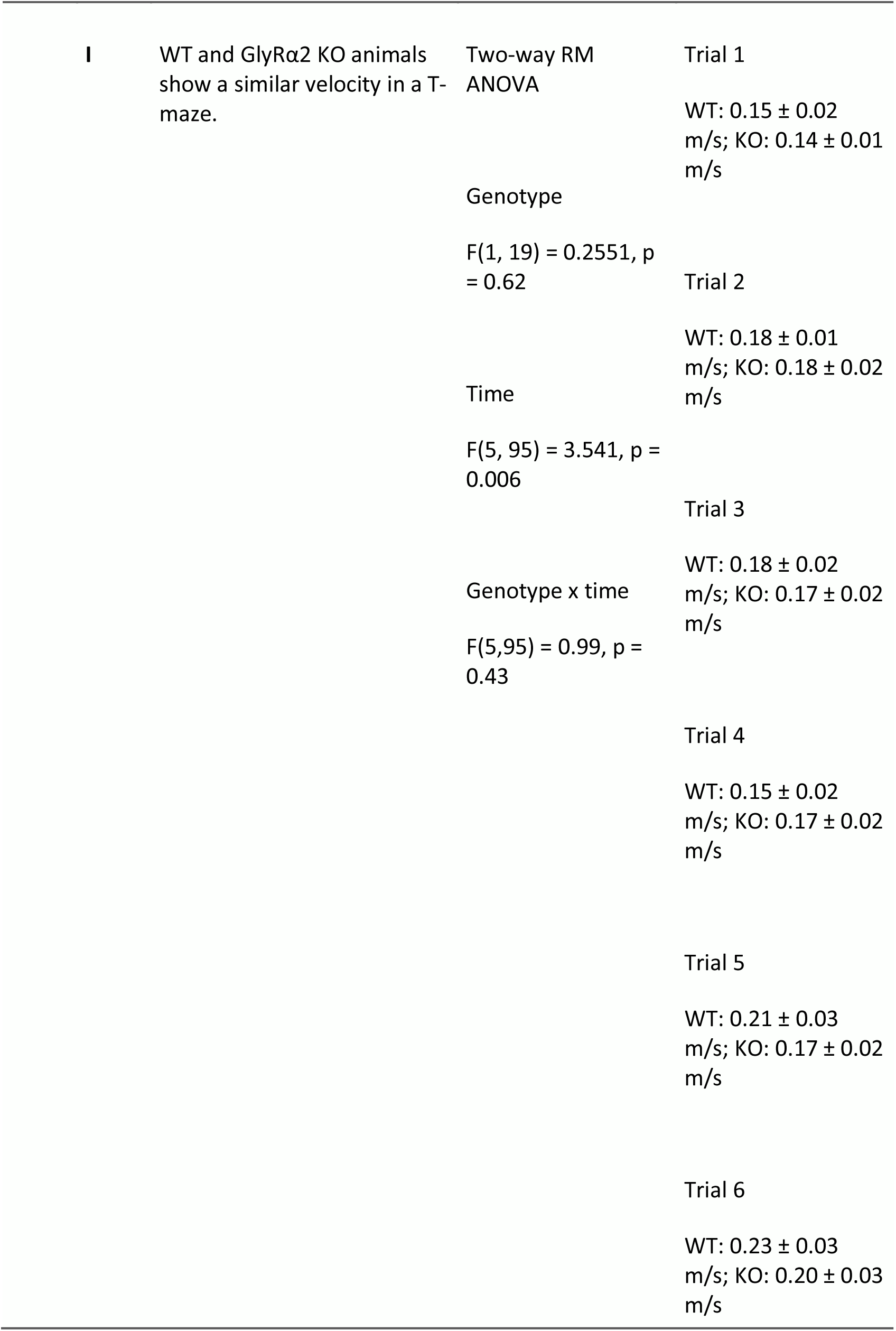
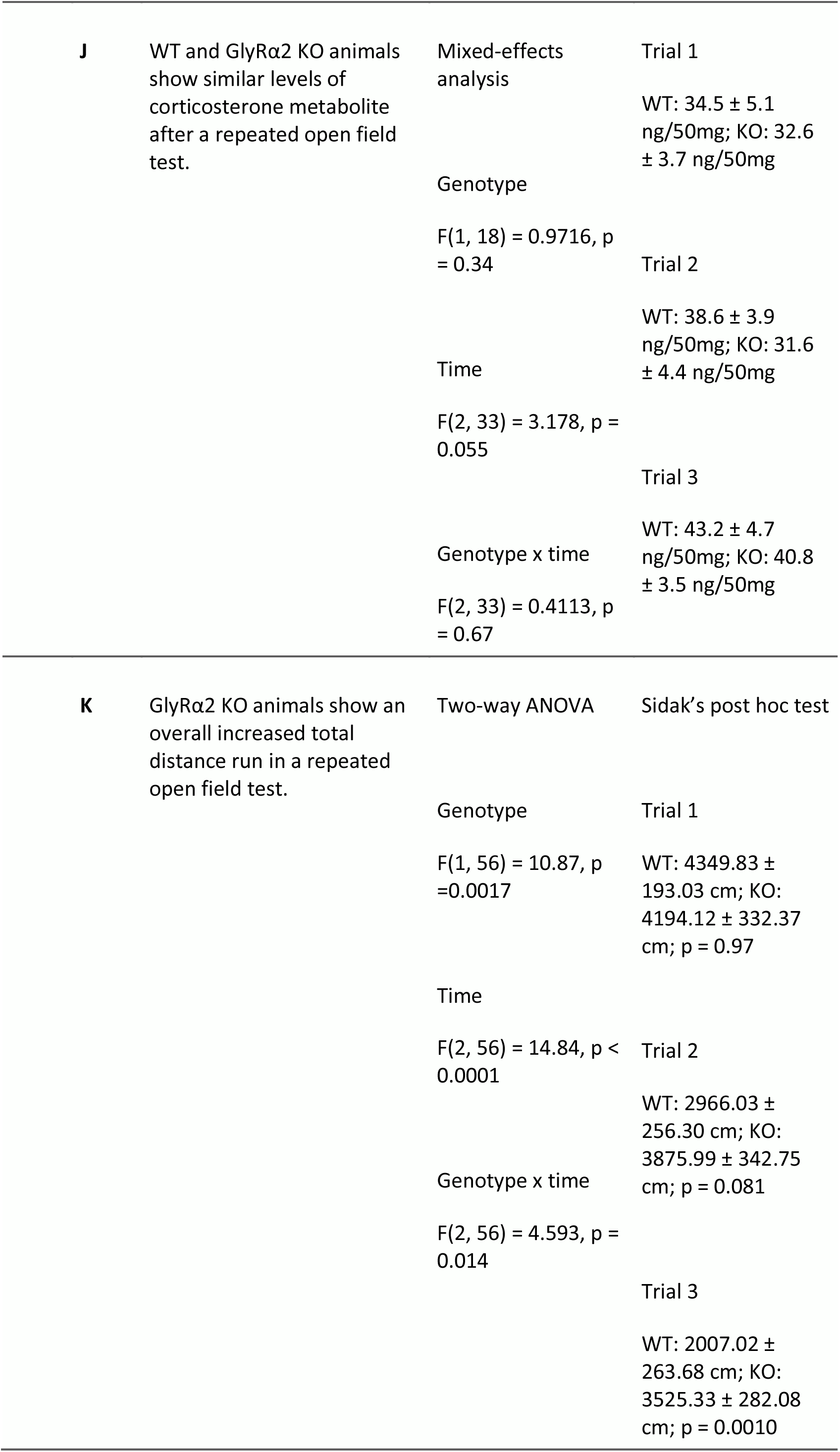
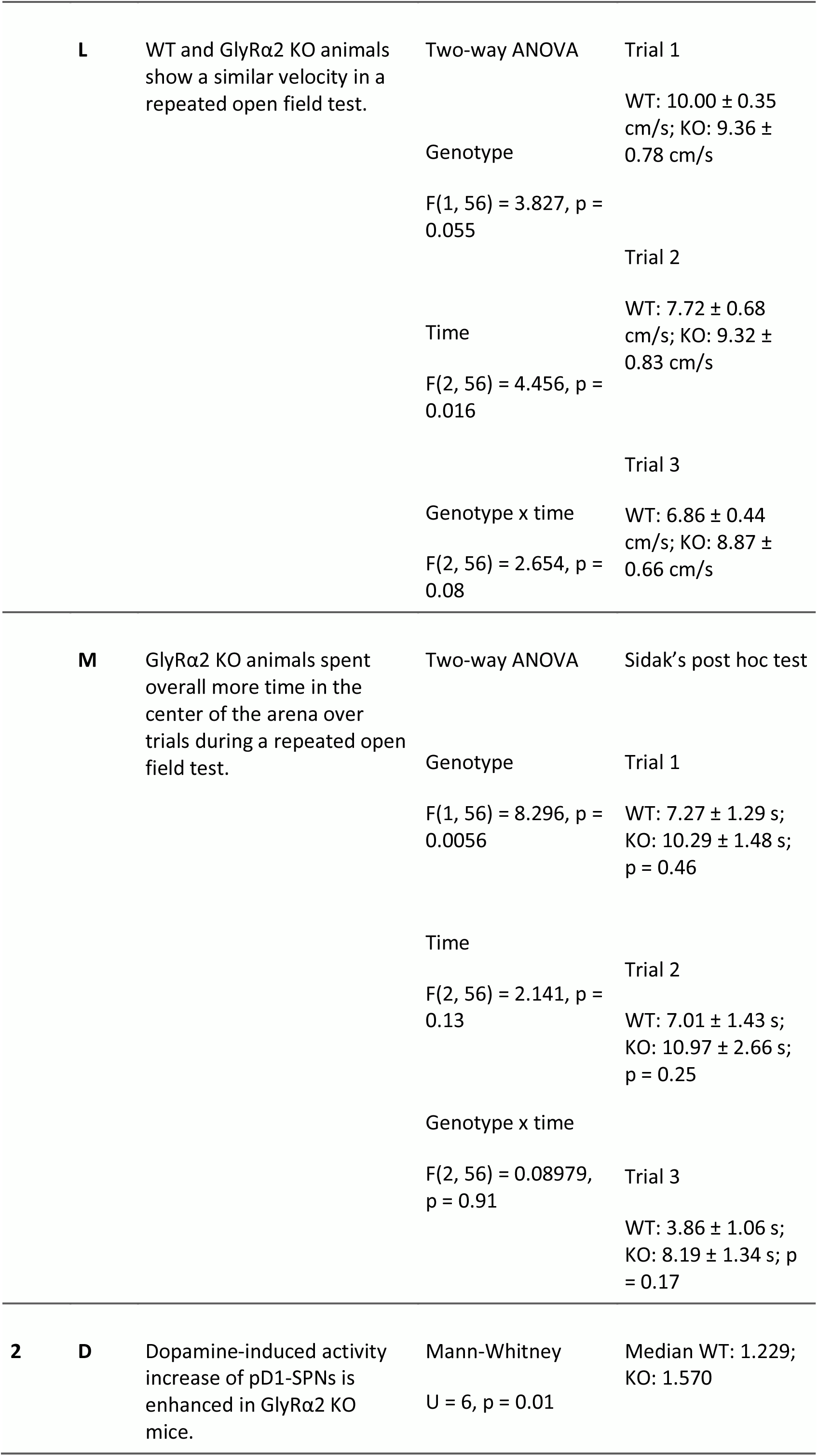
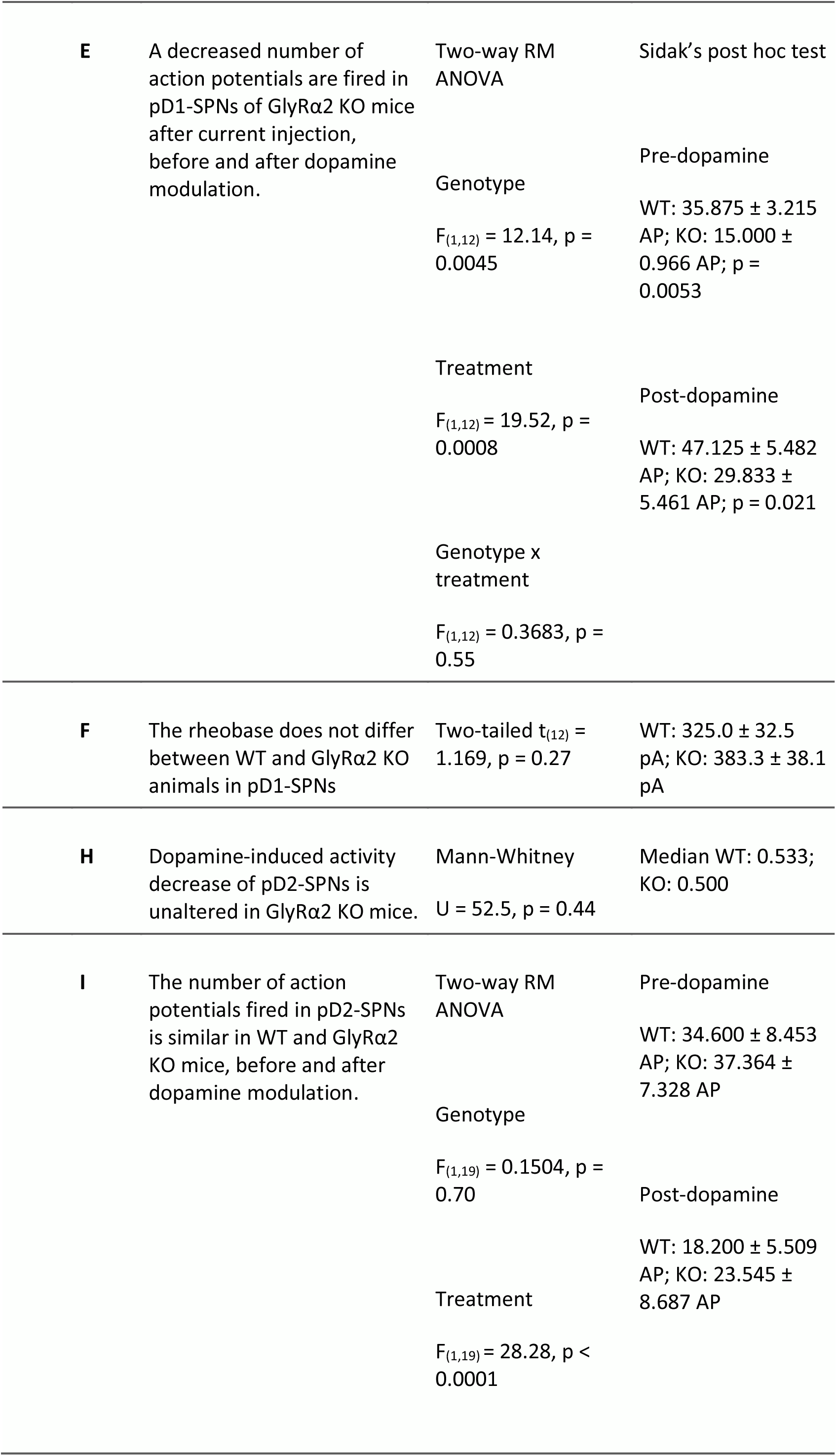
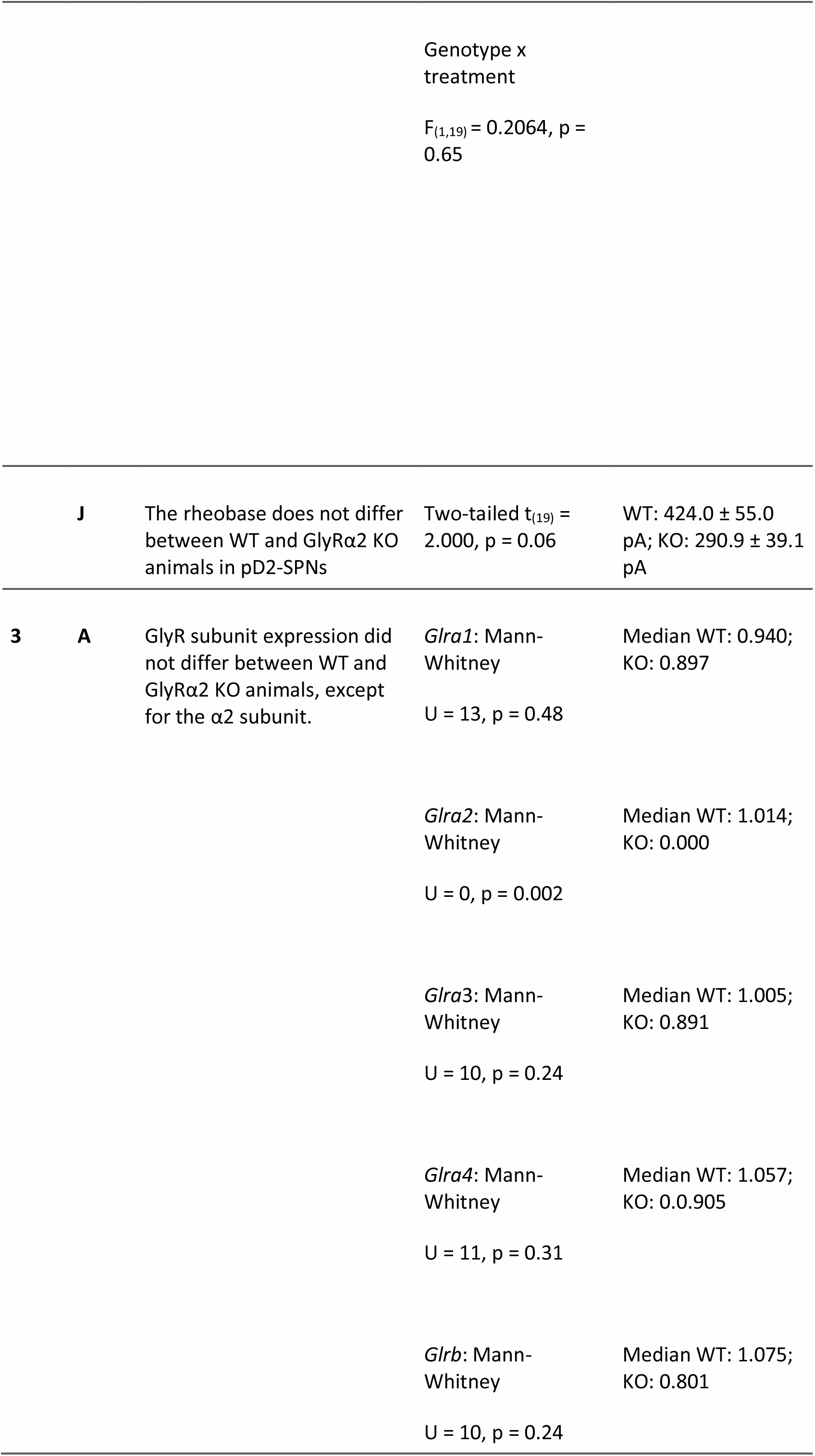
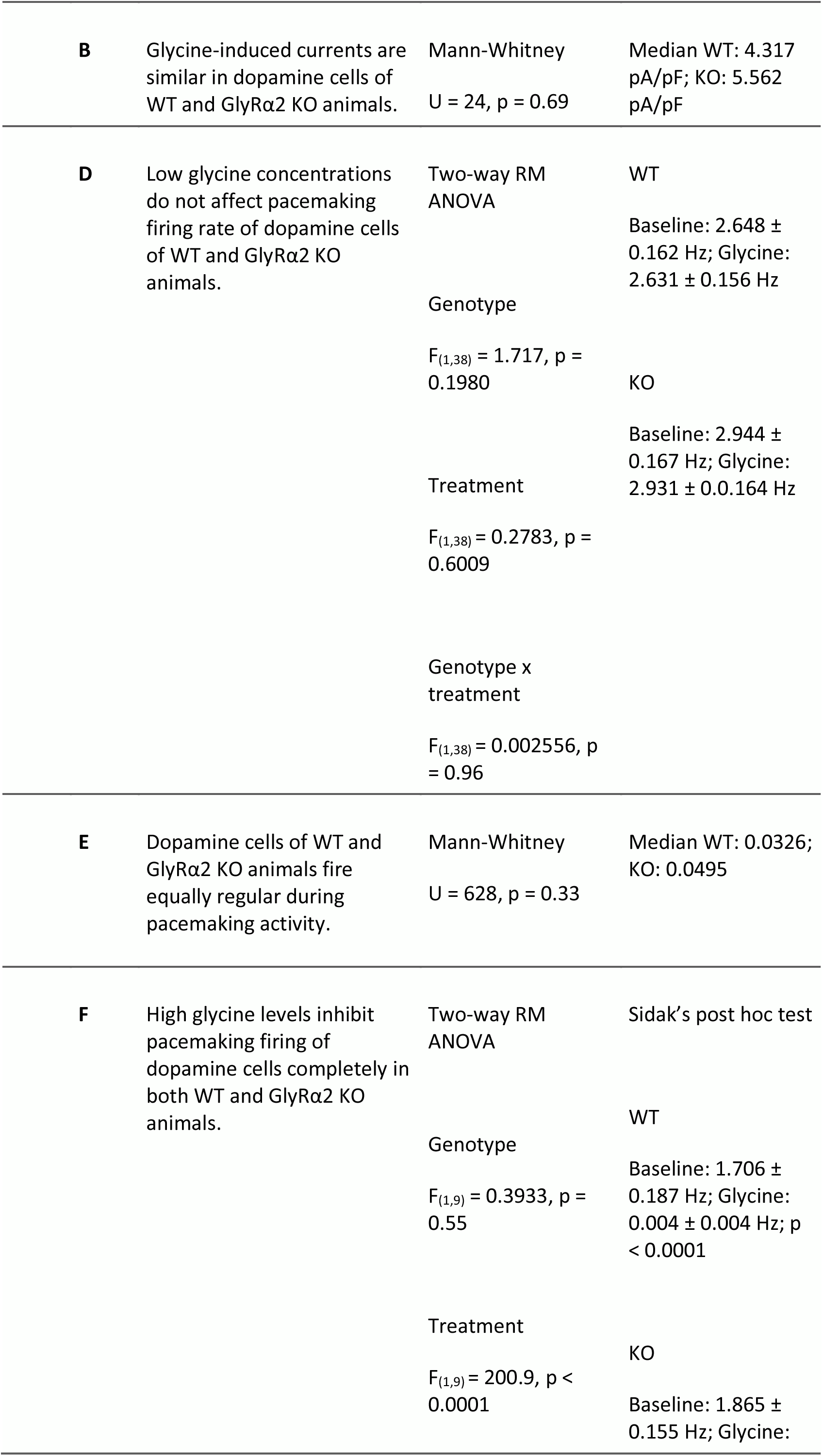
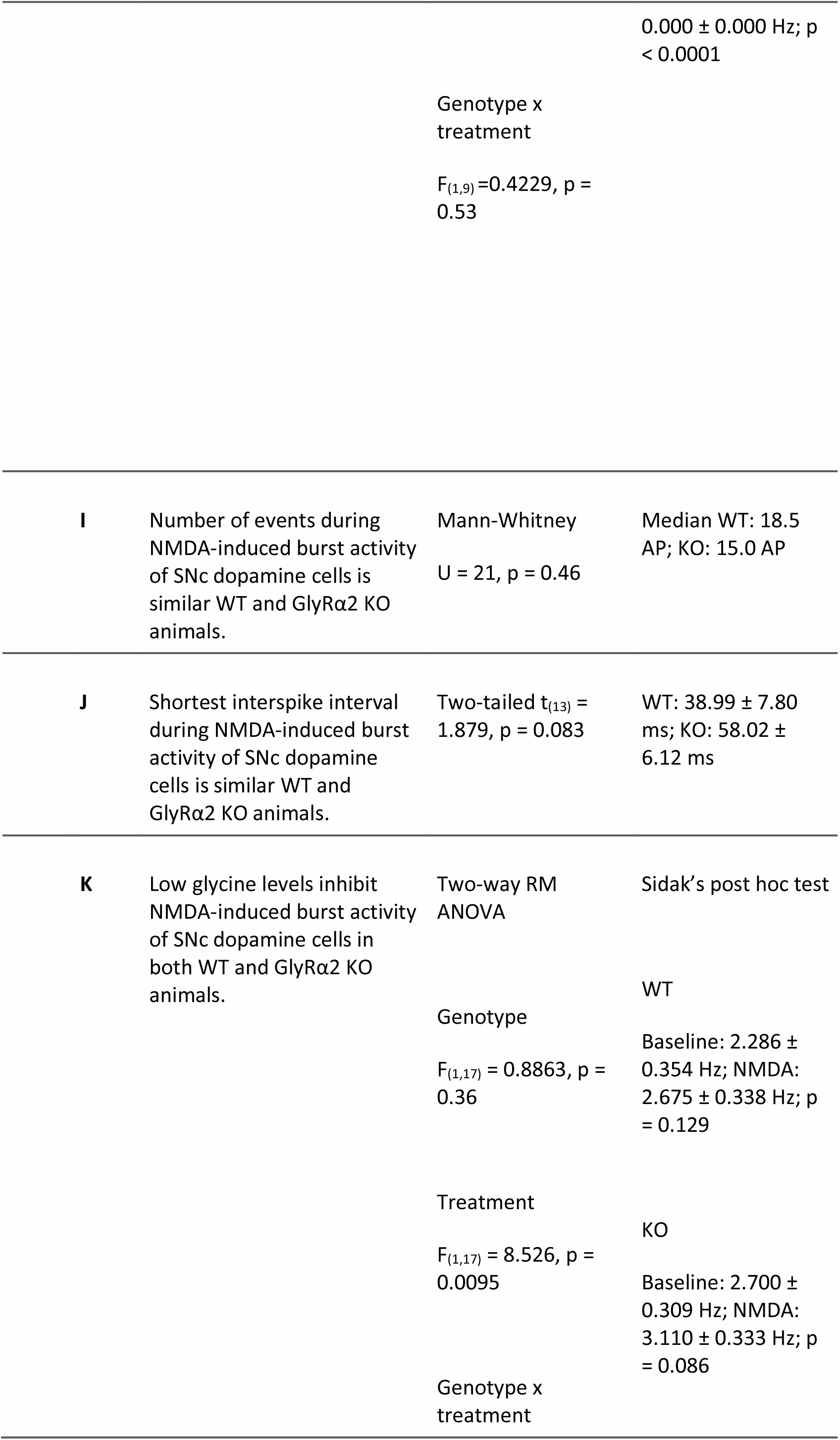
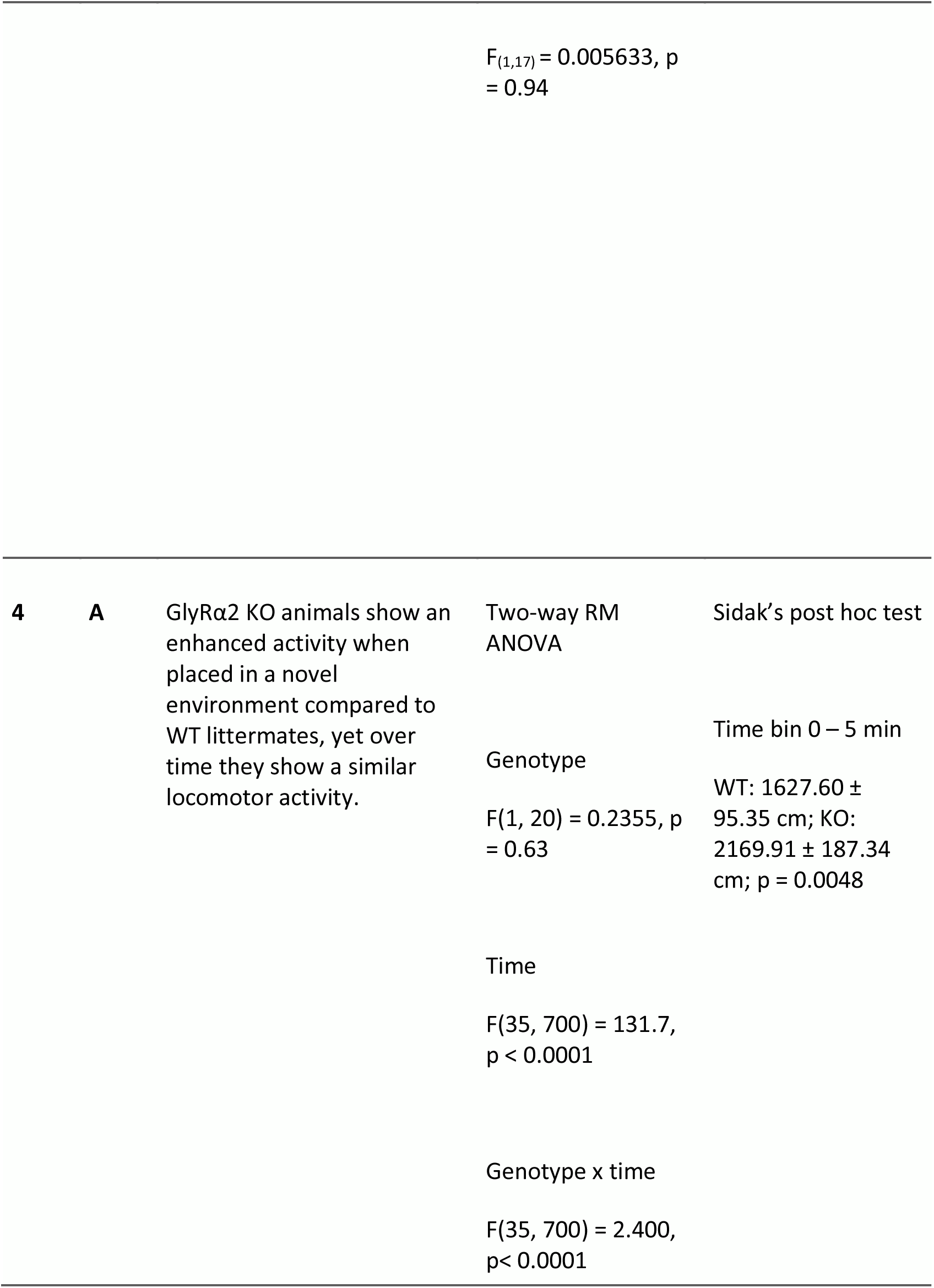
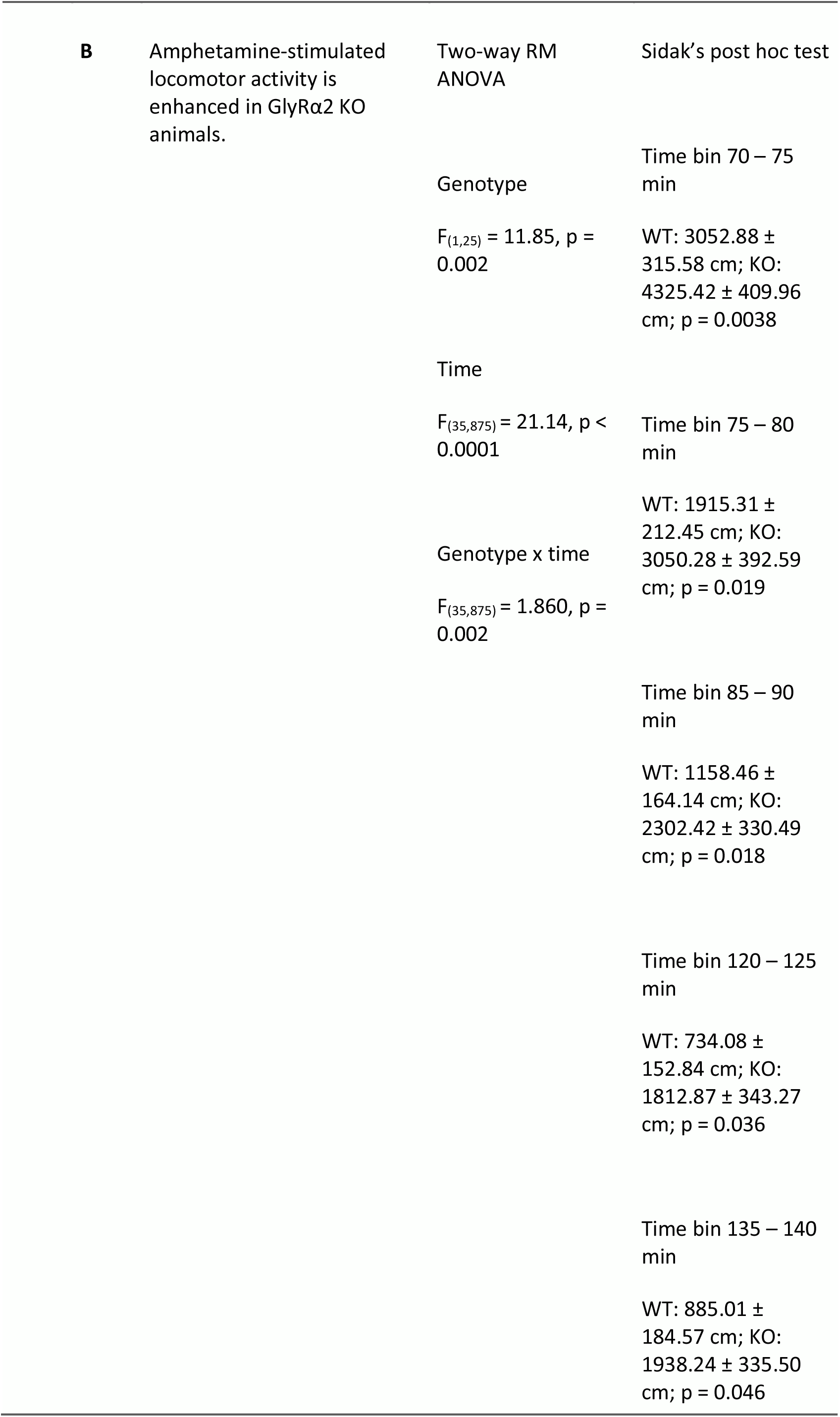
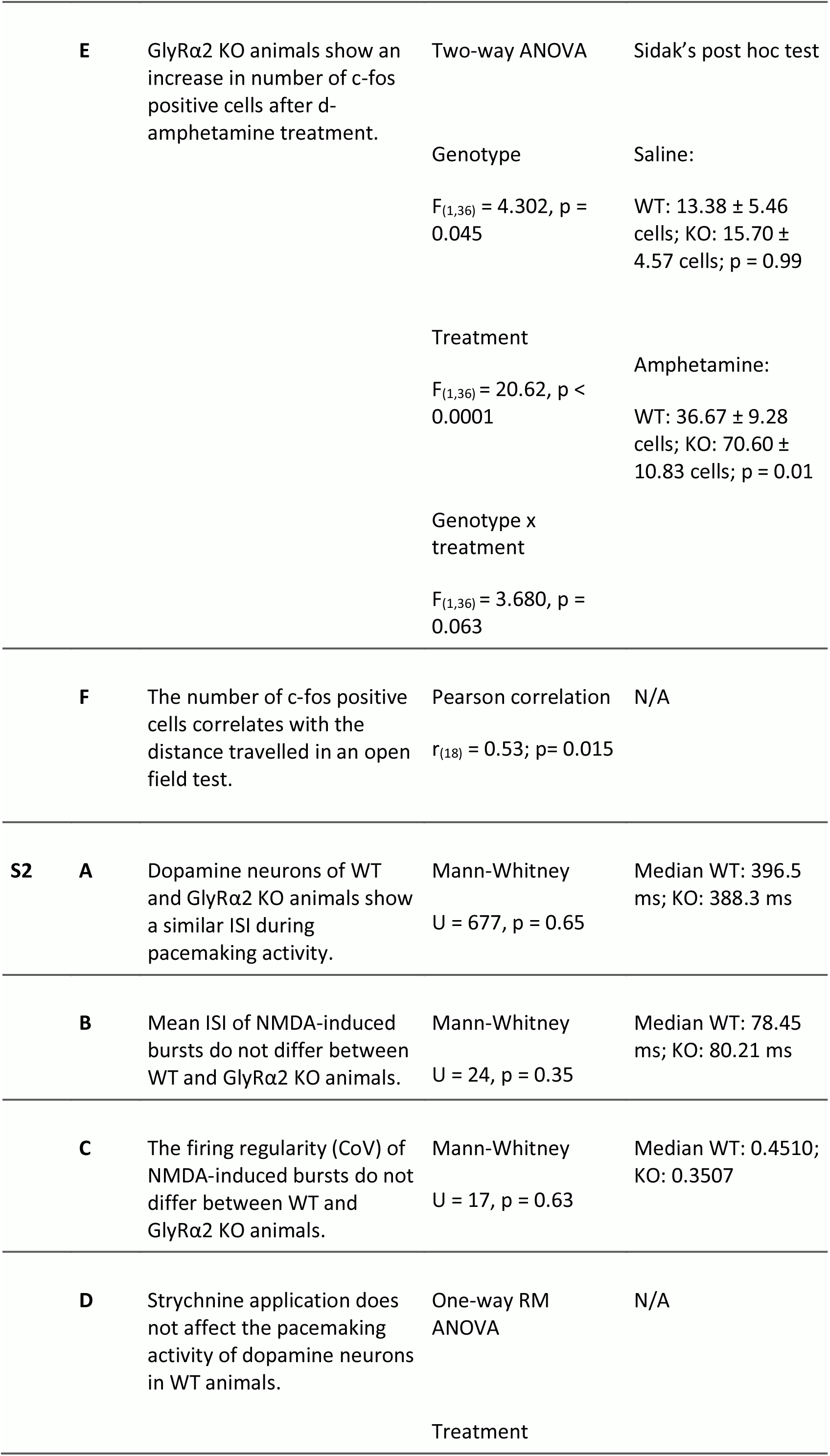
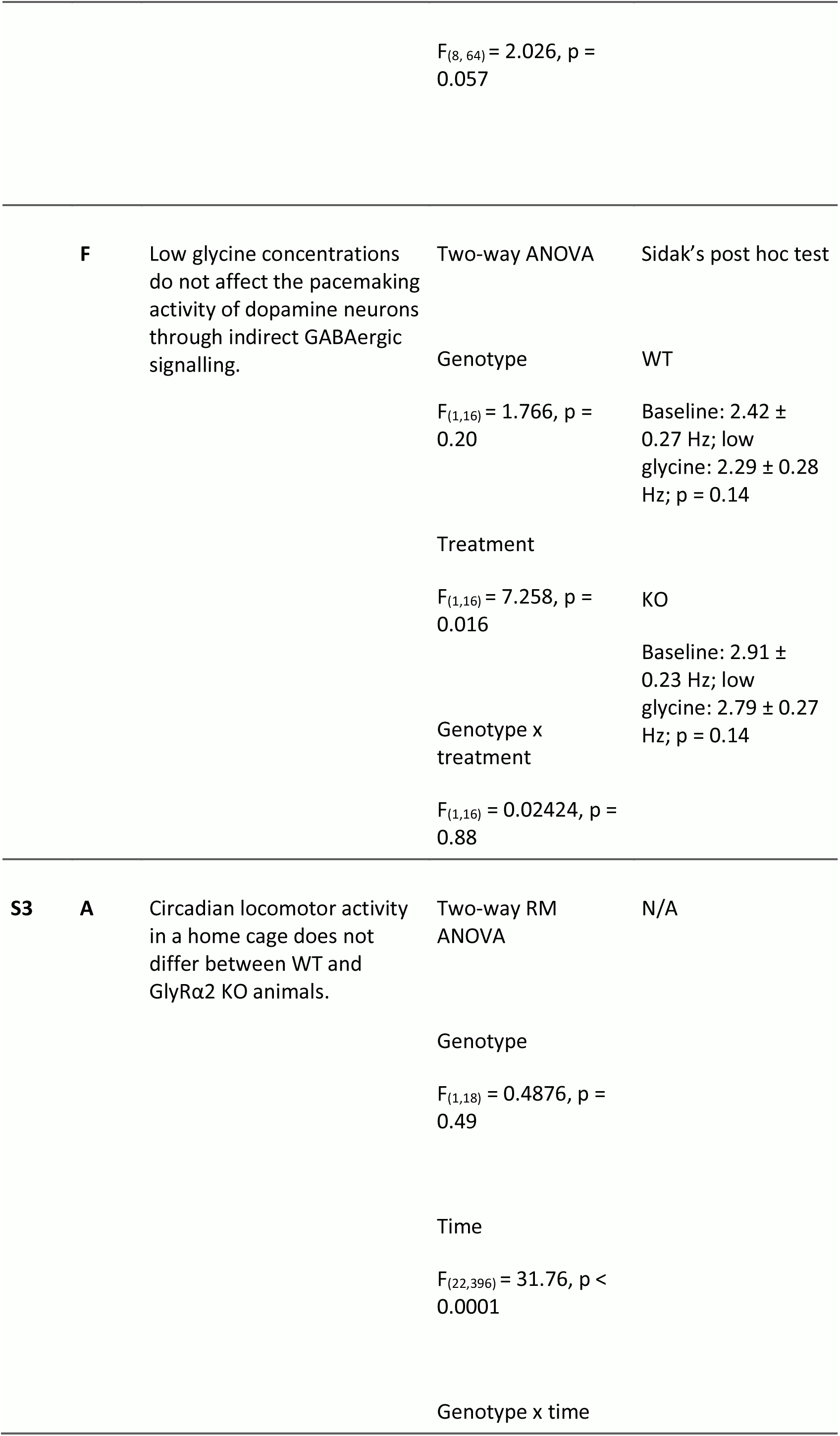
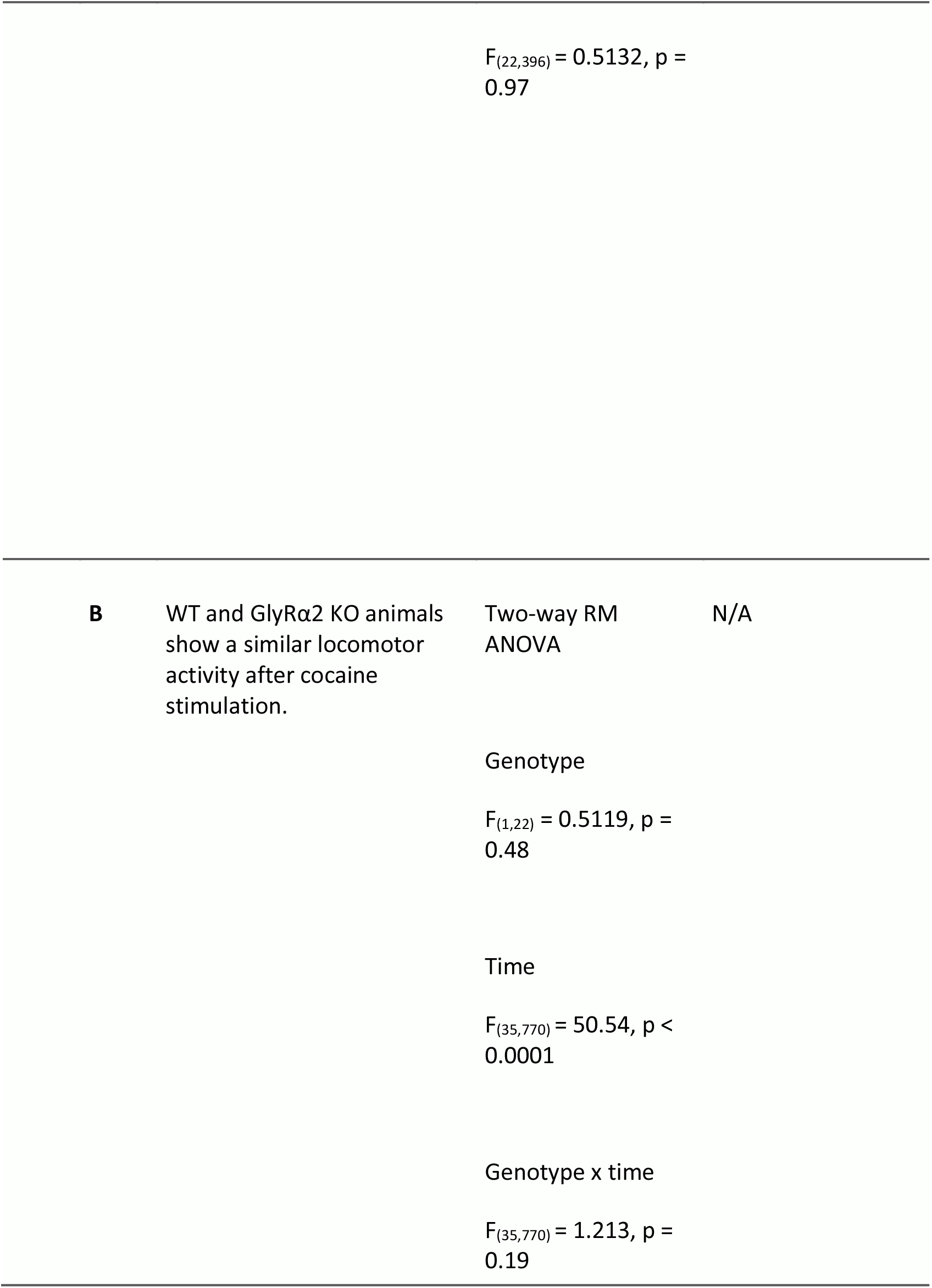
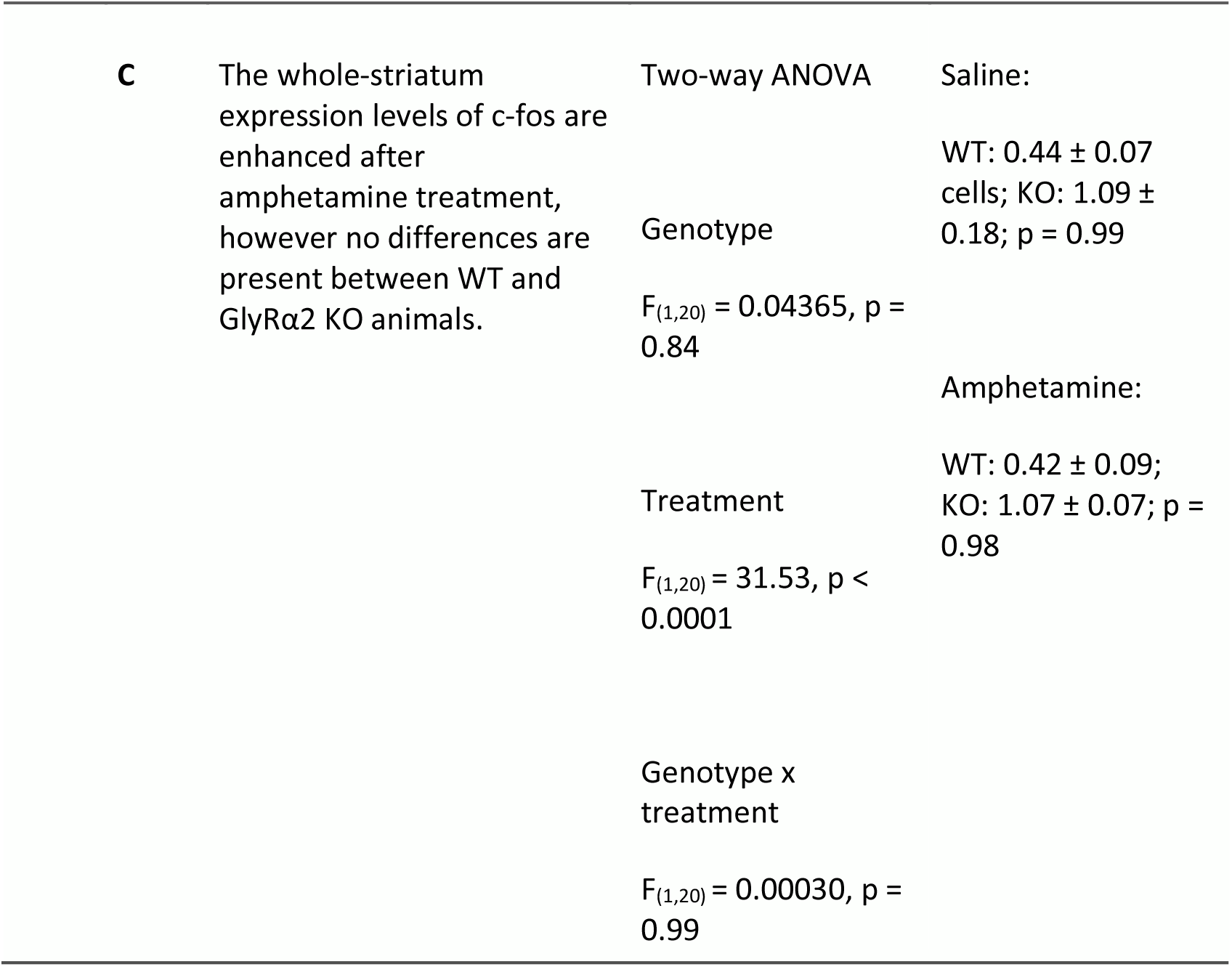

**Supplemental table 2.**

| Membrane resistance |  |  |  |
| --- | --- | --- | --- |
|  | WT (Mean ± SEM) | GlyRα2 KO (Mean ± SEM) | Statistical test |
| p-D1 SPNs | 55.8 ± 8.2 MOhm | 41.8 ± 2.6 MOhm | Mann-Whitney<br>U = 13, p = 0.18 |
| p-D2 SPNs | 42.7 ± 4.7 MOhm | 61.5 ± 6.7 MOhm | Mann-Whitney<br>U = 29, p = 0.07 |

**Supplemental table 3.**
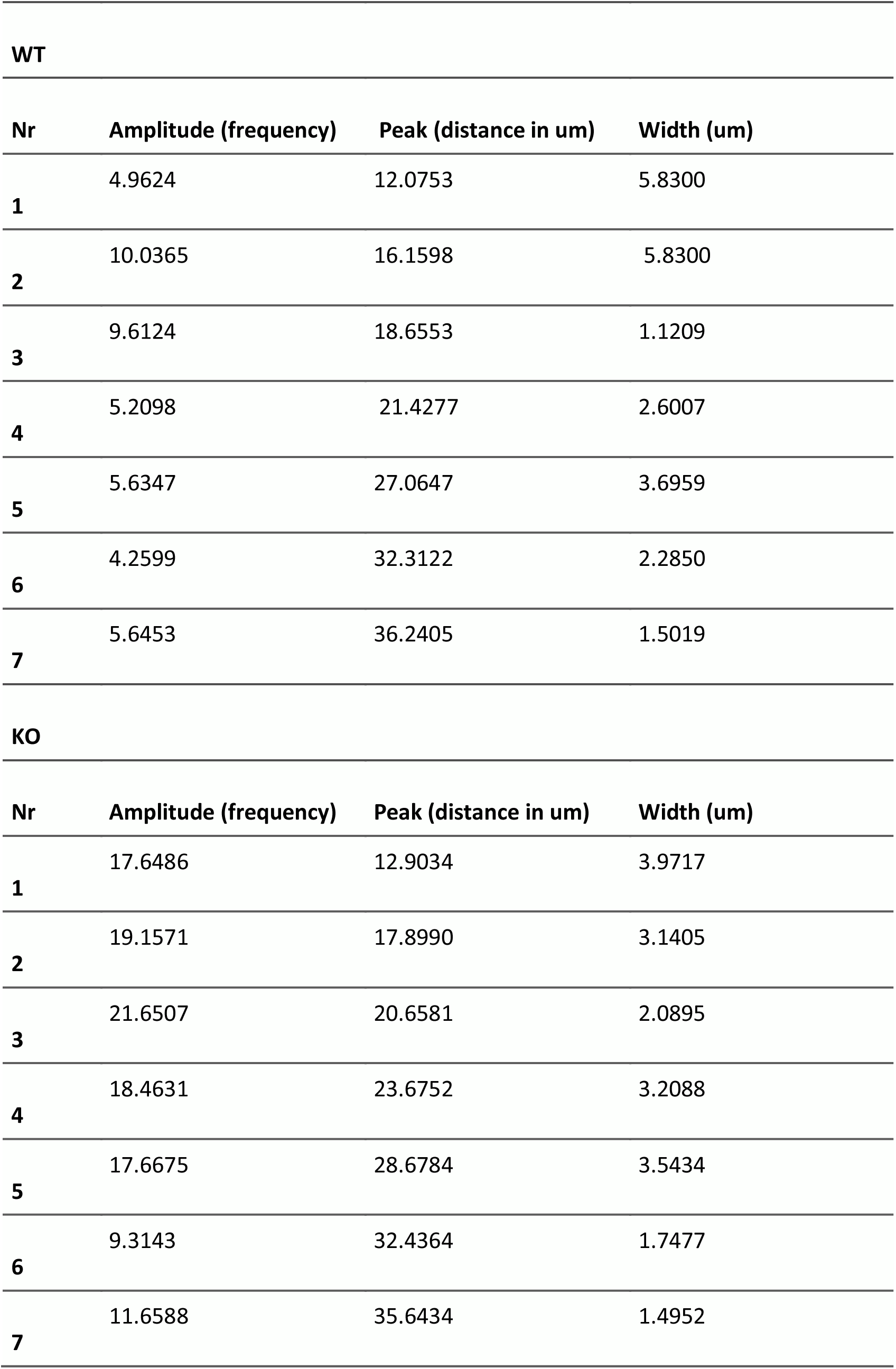

## Notes

### Competing Interest Statement

The authors have declared no competing interest.

### Summary of Updates

The manuscript has been revised after peer review.

